# The genomic landscape, causes, and consequences of extensive phylogenomic discordance in Old World mice and rats

**DOI:** 10.1101/2023.08.28.555178

**Authors:** Gregg W. C. Thomas, Jonathan J. Hughes, Tomohiro Kumon, Jacob S. Berv, C. Erik Nordgren, Michael Lampson, Mia Levine, Jeremy B. Searle, Jeffrey M. Good

## Abstract

A species tree is a central concept in evolutionary biology whereby a single branching phylogeny reflects relationships among species. However, the phylogenies of different genomic regions often differ from the species tree. Although tree discordance is often widespread in phylogenomic studies, we still lack a clear understanding of how variation in phylogenetic patterns is shaped by genome biology or the extent to which discordance may compromise comparative studies. We characterized patterns of phylogenomic discordance across the murine rodents (Old World mice and rats) – a large and ecologically diverse group that gave rise to the mouse and rat model systems. Combining new linked-read genome assemblies for seven murine species with eleven published rodent genomes, we first used ultra-conserved elements (UCEs) to infer a robust species tree. We then used whole genomes to examine finer-scale patterns of discordance and found that phylogenies built from proximate chromosomal regions had similar phylogenies. However, there was no relationship between tree similarity and local recombination rates in house mice, suggesting that genetic linkage influences phylogenetic patterns over deeper timescales. This signal may be independent of contemporary recombination landscapes. We also detected a strong influence of linked selection whereby purifying selection at UCEs led to less discordance, while genes experiencing positive selection showed more discordant and variable phylogenetic signals. Finally, we show that assuming a single species tree can result in high error rates when testing for positive selection under different models. Collectively, our results highlight the complex relationship between phylogenetic inference and genome biology and underscore how failure to account for this complexity can mislead comparative genomic studies.

## Introduction

Phylogenies are the unifying concept in understanding the evolution of species, traits, and genes. However, with the availability of extensive high-throughput sequencing data it has become apparent that evolutionary relationships between species may not be well represented by a single representative phylogeny (Edwards 2009; Hahn and Nakhleh 2016). While a dominant signal of bifurcating speciation may exist (*i.e.*, a species tree), phylogenetic signal that may disagree with species relationships can arise from ancestral polymorphisms (incomplete lineage sorting; ILS), gene flow through hybridization (introgression), and gene duplication and loss (Maddison 1997). The theoretical prediction of phylogenetic discordance has long been appreciated (Hudson 1983; Pamilo and Nei 1988; Maddison 1997; Rosenberg 2002), but empirical evidence now emphasizes just how extensive discordance can be among a set of species. For instance, studies of birds (Jarvis et al. 2014), seals (Lopes et al. 2021), tomatoes (Pease et al. 2016), and insects (Sun et al. 2021) have found that with extensive taxon sampling, highly supported species trees are rarely or never recovered in the underlying gene-trees. Whereas these examples highlight the prevalence of phylogenetic discordance across the tree of life, we still lack a clear understanding of how phylogenetic patterns are shaped by the details of genome biology or the extent to which discordance may compromise inferences from comparative studies that assume a singular species history.

From a practical perspective, failure to acknowledge and account for phylogenetic discordance could severely affect biological inference. Analyses of molecular evolution are usually performed on a gene-by-gene basis (Pond et al. 2005; Yang 2007; Hu et al. 2019; Kowalczyk et al. 2019), but it is still common practice to assume a single genome-wide species tree. For gene-based analyses, using the wrong tree may compromise inferences of positive directional selection, convergent evolution, and genome-wide inferences of correlated rate variation (Mendes et al. 2016). Phylogenetic discordance can also affect how continuous traits are reconstructed across phylogenies, as the genes that underly these traits may not follow the species history (Avise and Robinson 2008; Hahn and Nakhleh 2016; Mendes et al. 2018). In these instances, phylogenetic discordance may need to be characterized and incorporated into the experimental and analytical design. Alternatively, if a researcher’s primary questions are focused on reconstructing the evolutionary history of speciation (*i.e.*, the species tree), then phylogenetic discordance may obscure the true signal of speciation (Fontaine et al. 2015). In this case, knowledge about patterns of discordance across genomes could inform decisions about locus selection, data filtering, and model parameters during species tree reconstruction.

Given these considerations, a better understanding of the genomic context of phylogenetic discordance is warranted. Although often conceptualized primarily as a stochastic consequence of population history (Maddison 1997), patterns of phylogenetic discordance are likely to be non-random and structured across the genome, dependent on localized patterns of genetic drift, natural selection, recombination, and mutation. Discordance due to ILS ultimately depends on effective population sizes across the phylogeny (Pamilo and Nei 1988; Degnan and Rosenberg 2006) and, therefore, should covary with any process that influences local patterns of genetic diversity (*e.g*., linked negative or positive selection). Likewise, the potential for discordance due to introgression may be influenced by selection against incompatible alleles or, less often, positive selection for beneficial variants (Lewontin and Birch 1966; Jones et al. 2018). These sources of discordance, ILS and introgression, are expected to leave differing signals across the genomes of a sample of species that should allow us to test hypotheses about both the cause and the scale of phylogenetic discordance (Huson et al. 2005; Kulathinal et al. 2009; Green et al. 2010; Vanderpool et al. 2020). Yet the genomic context of phylogenetic discordance has remained elusive. For example, the potential for any of these processes to generate localized patterns of phylogenetic discordance depends on patterns of recombination (Hudson and Kaplan 1988) and recent simulation studies posit that phylogenies in the genome are expected to be correlated based on distance – the closer two regions are in the genome, the more history they share (McKenzie and Eaton 2020). However empirical studies have been inconclusive regarding the relationship between phylogenetic discordance and mammalian recombination rates, ranging from no correlation in great apes (Hobolth et al. 2007) to a weak correlation in house mice (White et al. 2009). However, if recombination rates evolve sufficiently quickly, long-term discordance measured over evolutionary timescales might be largely independent of contemporary recombination landscapes. To investigate the causes and consequences of phylogenetic discordance, we took advantage of genomic resources available for house mouse (*Mus musculus*). This rodent species is one of the most important mammalian model systems for biological and biomedical research and is embedded within a massive radiation of Old World rats and mice (Murinae). This ecologically diverse and species-rich group is comprised of over 600 species and makes up >10% of all mammalian species, and yet is only about 15 million years old. Despite this diversity and the power of evolution-guided functional and biomedical analysis, few other murine genomes have been sequenced outside of *Mus* and *Rattus*, with most efforts having been focused on sampling variation across closely related lineages of house mice. In the present work, we analyze new genome sequences for seven murine rodent species (*Mastomys natalensis*, *Hylomyscus alleni*, *Praomys delectorum*, *Rhabdomys dilectus*, *Grammoyms dolichurus*, *Otomoys typus*, and *Rhynchomys soricoides*) sampled from across this radiation. These new genomes are a powerful new resource for studying functional biology within rodents. However, along with increased taxonomic sampling, the presence of phylogenetic discordance poses challenges to rigorous molecular evolution analyses. We investigate these patterns of discordance in rodents here by combining these new genomes with previously sequenced rodent genomes and genomic resources from the *M. musculus* model system to study the phylogenetic relationships within Murinae as well as the landscape of discordance along rodent chromosomes. We first inferred a species tree for these and other sequenced rodent genomes, focusing on signals derived from ultra-conserved elements (UCEs) to promote broader comparisons across genomes of variable quality. We then use whole genome sequences, genetic maps, and annotation information from *M. musculus* to describe the genomic context of phylogenetic discordance at a broader taxonomic scale and evaluate several hypotheses linking discordance to genetic drift, natural selection, and recombination. Finally, we show how the use of a single species-tree impacts gene-level inferences from common molecular evolution tests for natural selection in these species.

## Methods

### Sample collection

As first reported in Kumon et al. (2021), we obtained frozen tissue samples from male individuals from the Museum of Vertebrate Zoology, Berkeley, CA (MVZ) and the Field Museum of Natural History, Chicago, IL (FMNH). *Hylomyscus alleni* (MVZ Mamm 196246) was collected in Cameroon in 2000, *Praomys delectorum* (MVZ Mamm 221157), *Mastomys natalensis* (MVZ Mamm 221054) and *Grammomys dolichurus* (MVZ Mamm 221001) were collected in Malawi in 2007, *Rhabdomys dilectus* (FMNH 192475) in Malawi in 2006, *Rhynchomys soricoides* (FMNH 198792) in The Philippines in 2008, and *Otomys typus* (FMNH 230007) in Ethiopia in 2015. Genome assembly sources are summarized in Table 1.

**Table 1:** All taxa whose genomes were included in this study, the source of the assembly, and the assembly level of each genome.

| **Taxon** | **Assembly source** | **Assembly level** | **No. UCEs** | **Used in genome-wide discordance analyses** |
| --- | --- | --- | --- | --- |
| *Apodemus speciosus* | GenBank: GCA_002335545.1 | Scaffolds | 2336 |  |
| *Apodemus sylvaticus* | GenBank: GCA_001305905.1 | Scaffolds | 2510 |  |
| *Arvicanthis niloticus* | GenBank: GCA_011762505.1 | Chromosomes | 2563 |  |
| *Grammomys dolichurus* | Kumon et al., 2021: De novo assembled | Chromosomes | 2395 | * |
| *Hylomyscus alleni* | Kumon et al., 2021: De novo assembled | Chromosomes | 2392 | * |
| *Mastomys natalensis* | Kumon et al., 2021: De novo assembled | Chromosomes | 2483 | * |
| *Mus caroli* | GenBank: GCA_900094665.2 | Chromosomes | 2584 |  |
| *Mus musculus* | GenBank: GRCm38.p6/mm10 | Chromosomes | 2294 | * |
| *Mus pahari* | GenBank: GCA_900095145.2 | Chromosomes | 2556 |  |
| *Mus spretus* | GenBank: GCA_001624865.1 | Chromosomes | 2578 |  |
| *Otomys typus* | This study: De novo assembled | Scaffolds | 2627 |  |
| *Phodopus sungorus* | Moore et al., 2022: De novo assembled | Chromosomes | 2633 |  |
| *Praomys delectorum* | Kumon et al., 2021: De novo assembled | Chromosomes | 2549 | * |
| *Rattus norvegicus* | GenBank: GCA_015227675.2 | Chromosomes | 2425 |  |
| *Rattus rattus* | GenBank: GCA_011064425.1 | Chromosomes | 2443 |  |
| *Rhabdomys dilectus* | Kumon et al., 2021: De novo assembled | Chromosomes | 2546 | * |
| *Rhombomys opimus* | GenBank: GCA_010120015.1 | Scaffolds | 2627 |  |
| *Rhynchomys soricoides* | Kumon et al., 2021: De novo assembled | Chromosomes | 2570 | * |

### Sequencing and assembly

We sequenced the seven new genomes in the Center for Applied Genomics at Children’s Hospital of Philadelphia. First, we extracted high molecular weight DNA following the protocol provided by 10XGenomics (CG000072 Rev B Sample Preparation Demonstrated Protocol, DNA Extraction from Fresh Frozen Tissue). We assessed the quality of the extracted DNA with CG00019 Rev B Sample Preparation Demonstrated Protocol, High Molecular Weight DNA QC, and all the samples had a mean length greater than 50kb, and high enough concentration to dilute to 1ng/mL for library preparation. We then used Chromium Genome Reagent Kits v2 from 10xGenomics to prepare libraries of 2x150 base reads, with read 1 constituting 10xBarcode (16 bp) + nmer (6bp) + genome sequence (128 bp) and read 2 constituting genome sequence (150 bp). The i7 index used 8bp sample index, and the i5 index was not used. We calculated sequencing depth based on a putative genome size of 3 Gb and coverage 56x, following 10XGenomics R&D recommendation, and then sequenced the libraries with Illumina HiSeq. We used the LongRanger v2.2.2 (Marks et al. 2019) wgs-basic pipeline to analyze demultiplexed FASTQ files. This pipeline gave general QC statistics related to the 10X barcoding and number of read pairs present in the FASTQ files. All sample FASTQs contained more than 688M read pairs and have acceptable barcode diversity/% on whitelist. We used Supernova to assemble *de novo* genomes. Genome sequences and assemblies were first reported in (Kumon et al. 2021), except for *O. typus*. While DNA extraction and sequencing on the 10x Genomics platform for *O. typus* is the same as described in (Kumon et al. 2021), the library quality for this sample was too low for chromosome level assembly. Here, we instead assembled it into scaffolds with the express purpose of obtaining UCEs for phylogenetic analysis. Adapters and low-quality bases were trimmed from the reads using illumiprocessor (Faircloth 2013), which makes use of functions from trimmomatic (Bolger et al. 2014). All cleaned reads were de novo assembled using ABySS 2.3.1 (Jackman et al. 2017) with a Bloom filter (Bloom 1970) de Bruijn graph. The final *O. typus* scaffold assembly was 2.14GB (N50=9,211; L50=64,014; E-size=12,790).

In parallel, for all samples excluding *O. typus*, we generated reference-based pseudo-assemblies with iterative mapping using pseudo-it (Sarver et al. 2017) to minimize reference bias in our genome-wide phylogenetic analyses and to maintain collinearity between assemblies. We used the *Mus musculus* (mm10) genome as the reference for our pseudo-assembly approach. In this version of pseudo-it, we have updated the software to call and insert indels into the pseudo-assembly (https://github.com/goodest-goodlab/pseudo-it). Briefly, pseudo-it maps reads from each sample to the reference genome with BWA (Li 2013), calls variants with GATK HaplotypeCaller (Poplin et al. 2018), and filters SNPs and indels and generates a consensus assembly with bcftools (Danecek et al. 2021). The process is repeated, each time using the previous iteration’s consensus assembly as the new reference genome to which reads are mapped. In total, we did 3 iterations of mapping for each sample.

### Ultraconserved element (UCE) retrieval

To reconstruct a broad phylogeny of murine rodents, we combined our seven recently sequenced genomes with nine publicly available genomes from other Old World mice and rats (subfamily Murinae) as well as the genomes of two non-murine rodents, the great gerbil (*Rhombomys opimus*; (Nilsson et al. 2020) and the Siberian hamster (*Phodopus sungorus*; (Moore et al. 2022) as outgroups. We extracted UCEs from each species, plus 1000 flanking bases from each side of the element using the protocols for harvesting loci from genomes and the *M. musculus* UCE probe set provided with phyluce v1.7.1 (Faircloth et al. 2012; Faircloth 2016). In total, we recovered 2,632 unique UCE loci, though not all UCE loci were found in all taxa (Table 1).

### UCE alignment

We brought the extracted UCE sequences for each species into a consistent orientation using MAFFT v7 (Katoh and Standley 2013) and then aligned them using FSA (Bradley et al. 2009) with the default settings. We trimmed UCE alignments with TrimAl (Capella-Gutierrez et al. 2009) with a gap threshold of 0.5 and otherwise default parameters. We performed alignment quality checks using AMAS (Borowiec 2016). We processed all alignments in parallel with GNU Parallel (Tange 2018).

### Species tree reconstruction from UCEs

We constructed a species-level rodent phylogeny with two approaches. First, using the alignments of all UCEs found in four or more taxa (2,632), we reconstructed a maximum-likelihood (ML) species tree with IQ-TREE v2.2.1 (Minh et al. 2020b). Each UCE alignment was concatenated and partitioned (Chernomor et al. 2016) such that optimal substitution models were inferred for individual UCE loci with ModelFinder (Kalyaanamoorthy et al. 2017). We also reconstructed individual gene trees for each UCE alignment. For all IQ-TREE runs (concatenated or individual loci), we assessed branch support with ultrafast bootstrap approximation (UFBoot) (Hoang et al. 2018) and the corrected approximate likelihood ratio test (SH-aLRT) (Guindon et al. 2010). We collapsed branches in each UCE tree exhibiting less than 10% approximated bootstrap support using the nw_ed function from Newick Utilities (Junier and Zdobnov 2010). We used these trees as input to the quartet summary method ASTRAL-III v5.7.8 (Zhang et al. 2018) to infer a species tree. We generated visualizations of phylogenies with R v4.1.1 (R Core Team 2021) using phytools v1.9-16 (Revell 2012) and the ggtree package v3.14 (Yu et al. 2017; Yu 2020) and its imported functions from ape v5.0 (Paradis and Schliep 2019) and treeio v1.16.2 (Wang et al. 2020).

We then used two methods to assess phylogenetic discordance across the reconstructed species tree. First, we calculated site and gene concordance factors (sCF and gCF) with IQ-TREE 2 (Minh et al. 2020a; Minh et al. 2020b) to assess levels of phylogenetic discordance between the inferred UCE trees and the concatenated species tree. gCF is calculated for each branch in the species tree as the proportion of UCE trees in which that branch also appears (Baum 2007). sCF represents the proportion of alignment sites concordant with a given species tree branch in a randomized subset of quartets of taxa (Minh et al. 2020a). We visualized gCF and sCF (Lanfear 2018) for each branch in each species tree using methods in R v4.3.0 (Lanfear 2018; R Core Team 2021). Next, we used PhyParts (Smith et al. 2015b) to identify topological conflict between the UCE trees and the species tree from ASTRAL-III. For this analysis, we rooted all trees with *Phodopus sungorus* as the outgroup using the nw_reroot function in the Newick Utilities (Junier and Zdobnov 2010) package and excluded 204 UCE trees that did not contain the outgroup.

### Divergence time estimation

We used IQ-TREE 2’s (Minh et al. 2020b) implementation of least square dating to estimate branch lengths of our species trees in units of absolute time (To et al. 2016). To improve divergence-time estimation, we used SortaDate (Smith et al. 2018) to identify a set of 100 UCE loci that exhibit highly clocklike behavior and minimized topological conflict with the concatenated species tree. We applied node age calibrations (Table 2) from Schenk et al. (2013) and Steppan and Schenk (2017), which in turn were sourced from fossil calibrations described on Paleobiology Database (2011). As *Rattus* is paraphyletic, the maximum age is taken from the earliest crown group fossil on Paleobiology Database (2011). In contrast, the estimated *Rattus* node age from Schenk et al (2013) was used as the minimum age. Branch lengths were resampled 100 times to produce confidence intervals. To return a single solution, least square dating typically requires that one calibration be fixed and not a range. We selected one calibration node (here, the branch leading to Murinae) and estimated dates across the tree when this node is set to its minimum, its maximum, and its midpoint ages. On the midpoint calibrated tree, we plot confidence intervals for each node representing the lowest minimum and highest maximum ages estimated across the three dating analyses.

**Table 2:** Prior node ages used in phylogenetic dating in millions of years before present.

| Clade | Minimum Age | Maximum Age | Citation |
| --- | --- | --- | --- |
| *Mus* | 5.3 | 7.2 | Steppan and Schenk, 2017 |
| *Apodemus* | 5.3 | 7.2 | Schenk et al., 2013 |
| *Rattus* | 2.4 | 3.6 | Steppan and Schenk, 2017 |
| Murinae | 12.1 | 14.05 | Schenk et al., 2013 |

### Genome window-based phylogenetic analysis

To assess the distribution of phylogenetic discordance across the rodent genome, we limited subsequent analyses to six of the newly sequenced genomes (*Mastomys natalensis*, *Hylomyscus alleni*, *Praomys delectorum*, *Rhabdomys dilectus*, *Grammomys dolichurus*, and *Rhynchomys soricoides*). *Otomys typus* was excluded from these analyses due to the inadequacy of the library outlined above. For these genomes, we generated pseudo-assemblies using pseudo-it and the mm10 reference genome to retain collinearity between genomes while minimizing reference bias.

Next, we partitioned the genomes into 10 kilobase (kb) windows based on the coordinates in the reference *M. musculus* genome (mm10; (Mouse Genome Sequencing et al. 2002)) using bedtools makewindows (Quinlan and Hall 2010). These coordinates were converted between the reference and the consensus sequence for each genome using liftOver (Hinrichs et al. 2006). Note that this method assumes both collinearity of all genomes and similar karyotypes (see Discussion). We then removed windows from the subsequent analyses if (1) 50% or more of the window overlapped with repeat regions from the *M. musculus* reference RepeatMasker (Smit et al. 2013- 2015) file downloaded from the UCSC Genome Browser’s table browser (Hinrichs et al. 2006) or (2) 50% or more of the window contained missing data in 3 or more samples. Overlaps with repeat regions were determined with bedtools coverage (Quinlan and Hall 2010). We then aligned the 10kb windows with MAFFT (Katoh and Standley 2013), trimmed alignments with trimAl (Capella-Gutierrez et al. 2009), and inferred phylogenies for each with IQ-TREE 2 (Minh et al. 2020b) which uses ModelFinder to determine the best substitution model for each window (Kalyaanamoorthy et al. 2017). To assess patterns of tree similarity between windows on the same chromosome, we used the weighted Robinson-Foulds (wRF) (Robinson and Foulds 1981; Böcker et al. 2013) distance measure implemented in the phangorn library (Schliep 2011) in R (R Core Team 2021). This measure compares two trees by finding clades or splits present in one tree but not the other. The weighted version of this measure increments a score by the missing branch length in each tree for each mismatch and additionally the differences in branch length between the co-occurring branches in both trees, allowing us to capture differences in branch length even when topology does not differ (Robinson and Foulds 1979). Consequently, the resulting measure of wRF is in units of branch length, in our case for maximum likelihood trees this is expected number of substitutions per site. We compared wRF between trees from windows on the same chromosome to characterize (1) heterogeneity in patterns discordance along the chromosome and (2) whether tree similarity is correlated with distance between windows. For the second question, we sampled every window on a chromosome at increasing distance (in 10kb windows) until the distribution of wRF scores for all pairs of windows at that distance was not significantly different (Wilcox test, p > 0.01) than that of a sample of 12,000 measures of wRF between randomly selected trees on different chromosomes. We selected 12,000 as the random sample size because it roughly matched the number of windows on the largest chromosome (chromosome 1, n = 12,113). We used Snakemake 7 (Mölder et al. 2021) to compute window alignments and trees in parallel.

### Whole genome alignment between mouse and rat

We used minimap2 (Li 2018) to align the mouse (mm10) and rat (rnor6) (Gibbs et al. 2004) genomes to assess the impact of structural variation that spans the divergence of our subset of species used to in the discordance analyses. We downloaded the rat reference genome (rnor6) from the UCSC genome browser and for both genomes removed the Y chromosome and all smaller unplaced scaffolds, leaving only the X chromosome and autosomes 1-19 for mouse and 1-20 for rat. We then used minimap2 in whole genome alignment mode (-x asm20) to generate a pairwise alignment file from which we calculated alignment segment sizes and the distances between alignment segments. We visualized the alignment as a dot plot using the pafr package in R (https://github.com/dwinter/pafr).

### Recombination rate and functional annotation

We retrieved 10,205 genetic markers generated from a large heterogenous stock of outbred mice (Shifman et al. 2006; Cox et al. 2009) to assess whether phylogenetic discordance along chromosomes is correlated with mouse recombination rates. We converted the physical coordinates of these markers from build 37 (mm9) to build 38 (mm10) of the *M. musculus* genome using liftOver (Hinrichs et al. 2006). We then partitioned the markers into 5Mb windows and estimated local recombination rates in each window. Estimated recombination rates were defined as the slope of the correlation between the location on the genetic map and the location on the physical map of the *M. musculus* genome for all markers in the window (White et al. 2009; Kartje et al. 2020). Within each 5Mb window, we calculated wRF distances between the first 10kb window and every other 10kb window.

We also compared the chromosome-wide wRF distances to those based on phylogenies from regions around several types of adjacent to genomic features. We retrieved coordinates from 25,753 protein coding genes annotated in *M. musculus* from Ensembl (release 99; (Cunningham et al. 2022)), all 3,129 UCEs from the *M. musculus* UCE probe set provided with PHYLUCE (Faircloth et al. 2012; Faircloth 2016), and 9,865 recombination hotspots from Smagulova et al. (2011). The recombination hotspot coordinates were converted between build 37 and build 38 using the liftOver tool (Hinrichs et al. 2006). For each feature, the starting window was the 10kb window containing the feature’s midpoint coordinate. We then calculated wRF between this window and all windows within 5Mb in either direction and for each chromosome compared the slope and wRF distance of windows adjacent to the feature with the same metrics for the whole chromosome. We compared distributions of these measures for each genomic feature with an ANOVA (aov(feature.measure ∼ feature.label)) followed by Tukey’s range test (TukeyHSD(anova.result)) to assess differences in means, as implemented in R v4.1.1 (R Core Team 2021).

### Molecular evolution

To test how tree misspecification affects common model-based analyses of molecular evolution, we retrieved 22,261 coding sequences from *M. musculus* using the longest coding transcript of each gene. Coding coordinates from the *M. musculus* coding sequences were transposed to the new assemblies via liftOver (Hinrichs et al. 2006) and sequences retrieved with bedtools getfasta (Quinlan and Hall 2010). Because some regions are too diverged, some genes are unable to liftOver for some samples so in total we recovered 17,216 genes present in all 7 species. We then used MACSE to re-align these coding regions to account for possible frameshifts and stop codons (Ranwez et al. 2018). Before alignment we trimmed non-homologous regions from each ortholog using the trimNonHomologousFragments sub-program within MACSE. Then we aligned the orthologs using alignSequences and trimmed the aligned sequences with trimAlignment to remove unaligned flanking regions. Finally, we manually filtered all alignments based on the following: We found that 3,368 alignments have one or more sequence removed during filtering for gappy sites, 3,132 alignments have a premature stop codon in at least one species, 1,571 alignments have only 3 or fewer unique sequences among the 7 species, and 78 alignments are shorter than 100bp. We removed these alignments from all subsequent analyses, resulting in 12,559 total alignments for tree reconstruction and inference of selection. Note that some alignments were filtered in multiple of the listed categories.

We then used IQ-TREE 2 (Minh et al. 2020b) to reconstruct a single species tree from concatenation of all gene alignments, as well as gene-trees for each individual alignment. This species tree from coding regions matches the topologies of these species inferred by concatenation of UCEs in the previous section. Next we ran several tests that use both coding alignments and a tree to infer positive selection: PAML’s M1a vs. M2a test (Yang 2007), HyPhy’s aBSREL model (Smith et al. 2015a), and HyPhy’s BUSTED model (Murrell et al. 2015). We ran each test twice on each gene, once using the species tree derived from concatenated data, and once using the gene tree inferred from the alignment of that gene only. For the HyPhy models, no target branch was selected, meaning all branches in the input phylogeny were tested.

The end point of each of each of these three tests is a p-value, which lets us assess whether a model that allows for positively selected sites fits better than a model that does not. For M1a vs. M2a, we obtained the p-value manually by first performing a likelihood ratio test to determine genes under selection by calculating 2 ∗ (𝑙𝑙𝑙𝑙𝑙𝑙 𝑀𝑀1𝑎𝑎 – 𝑙𝑙𝑙𝑙𝑙𝑙 𝑀𝑀2𝑎𝑎). The p-value of this likelihood ratio is then retrieved from a one-tailed chi-square distribution with 2 degrees of freedom (Yang 2007). For BUSTED and aBSREL, p-values are computed automatically during the test using similar likelihood ratios. For the M1a vs. M2a and BUSTED tests, a single p-value is computed for each gene. P-values were adjusted by correcting for false discovery rates (Benjamini and Hochberg 1995; Yekutieli and Benjamini 1999) using the “fdr” method in the p.adjust() function in R (R Core Team 2021) and we categorized a gene as being positively selected if its adjusted p-value was < 0.01. For the aBSREL test, a p-value is generated for each branch in the input gene tree. aBSREL corrects for multiple testing internally across branches using the Holm-Bonferroni procedure (Holm 1979; Pond et al. 2005). We further correct the p-values across genes with the Bonferroni method and classify a gene as having experienced positive selection if one or more branches has a p-value < 0.01 after all corrections. We used Snakemake 7 (Mölder et al. 2021) to compute coding alignments, trees, and selection tests in parallel.

## Results

### Murine species tree

We sequenced and assembled genomes from seven murine species that represent some of the first non-*Mus* or *Rattus* genomes to be sequenced from the large radiation of Old World mice and rats. We combined our new genomes with nine publicly available murine genomes as well as the giant gerbil, *Rhombomys opimus* (Nilsson et al. 2020) and the Siberian dwarf hamster, *Phodopus sungorus* (Moore et al. 2022). Using a concatenated dataset of 2,632 aligned ultra conserved elements (UCEs) we inferred a species tree (Fig. 1) that recovered the same relationships as previous reconstructions of Murinae using a small number of loci (Lecompte et al. 2008; Steppan and Schenk 2017). We also find that a species tree inferred from a quartet-based summary of the gene trees of all 2,632 UCEs is identical to one inferred from concatenation (Fig. S1).

**Figure 1:**
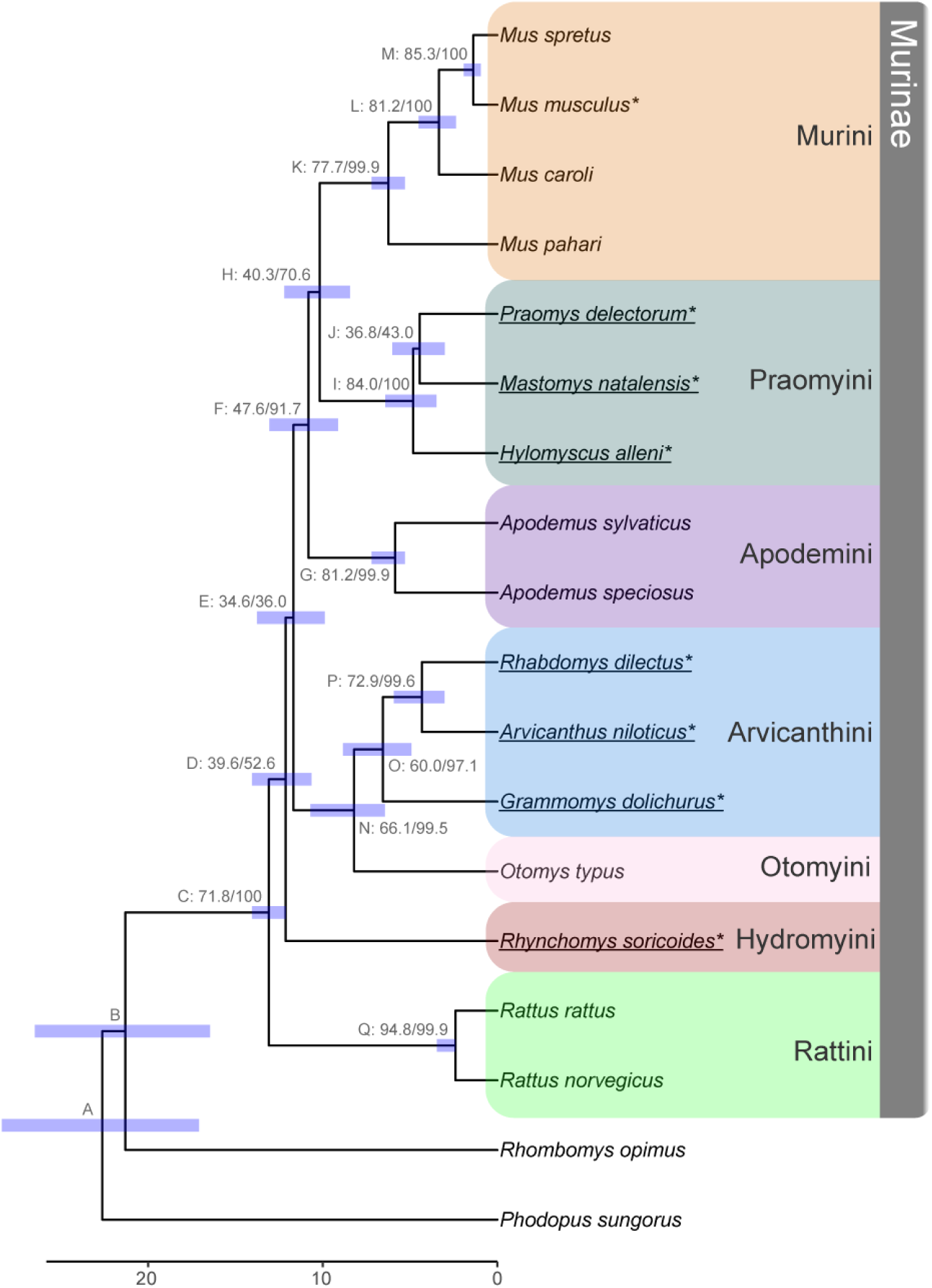
Species trees inferred from concatenation of ultra-conserved elements (UCEs) from 18 rodent species. Internal nodes are labeled by a letter identifier referenced in the text and site and gene concordance factors (*i.e.*, Label: sCF/gCF) as well as a bar indicating the confidence interval for divergence time estimation. Ultrafast bootstrap/SH-aLRT values were all 100. Bottom scale represents time in millions of years before present. Fossil calibrations are described in Tables 2 and S2, with node C used as a fixed calibration point. Tribes within sub-family Murinae are highlighted on the right following the classifications used by Lecompte et al. (2008). New genomes presented in this study are underlined, and genomes used for the genome-wide phylogenetic discordance analysis are indicated with an asterisk.

While bootstrap and SH-aLRT values provided high support to our inferred species trees (Fig. 1), we found evidence for discordance across individual UCE phylogenies. The five shortest branches in the concatenated tree had a site concordance factor (sCF) of less than 50%, suggesting that alternate resolutions of the quartet had equivocal support (Fig. S2). Gene concordance factors (gCF) for each branch in the species tree were on aggregate much higher, with all but four branches supported by almost every gene tree in the analysis and with the lowest values likely being driven by a several short internal branches (Fig. S2). This pattern is recapitulated under a coalescent model (Figs. S1 and S3). At the two most discordant nodes (E and J in Fig. 1), the recovered topology was supported by approximately one third of all gene trees.

We estimated divergence times for the inferred concatenated phylogeny (Fig. 1; Table S1) using four fossil calibration points (Table 2). The murid and cricetid groups had an estimated divergence time of 22.62 Ma (node A in Fig. 1) followed by the Murinae and the Gerbillinae at 21.30 Ma (B), albeit with wide confidence intervals in both cases. With the ancestral Murinae node. (C) fixed for calibration, Hydromyini arose at 12.12 Ma (D) and was followed by Otomyini and Arvicanthini at 11.67 Ma (E). The remaining Murine tribes evolved in rapid succession, with Apodemini diverging at 10.82 Ma (F) and Murini and Praomyini splitting at 10.08 Ma (H). The *Rattus* node, which was fossil calibrated, was recovered at the very youngest end of the calibration range.

### The landscape of phylogenetic discordance along murine genomes

Next, we described the genomic landscape of phylogenetic discordance in murines and its relationship to local genomic features. To do this, we analyzed genome-wide phylogenetic histories of six newly sequenced murine rodent genomes (excluding *Otomys typis* due to low assembly quality; see Fig. 1) and the well-annotated *M. musculus* reference genome. Using the coordinate system from the *M. musculus* genome we partitioned and aligned 263,389 non-overlapping 10 kb windows from these seven species (Table 1).

After we filtered windows in repetitive regions or with low phylogenetic signal, we recovered 163,765 phylogenies with an average of 616 informative sites per window (Fig. S4). We found that phylogenetic discordance was pervasive within and between chromosomes. We inferred 597 of the 945 possible unique rooted topologies among 6 species (when specifying *R. soricoides* as the outgroup) across all chromosomes (Table 3 shows the most common topologies recovered). The number of unique topologies per chromosome ranged from 75 to 218 with an average of 141 (Table 4). We ranked the recovered topologies by count per chromosome and found that just four different topologies were ranked in the top three in at least one chromosome. (Fig 2A; Table 3) and only nine are present at a frequency above 1%. Among these, the top three topologies only differ in the ordering of the clade containing *Hylomyscus alleni, Mastomys natalensis,* and *Praomys delectorum* (HMP clade) with the rest of the tree being constant. This clade also showed the second lowest concordance in the species tree inferred from UCEs (Fig. 1, node J) These three species trees each comprise roughly 14% of all recovered topologies (Fig. 2), though interestingly of the three, it is the least common one that matches the topology recovered via concatenation of all coding regions and the species tree recovered from UCEs (Fig. 1). However, this topology was only inferred for 13.1% of windows. That is, the robustly and consistently inferred species tree did not match the evolutionary relationships inferred for nearly 90% of the genome.

**Figure 2:**
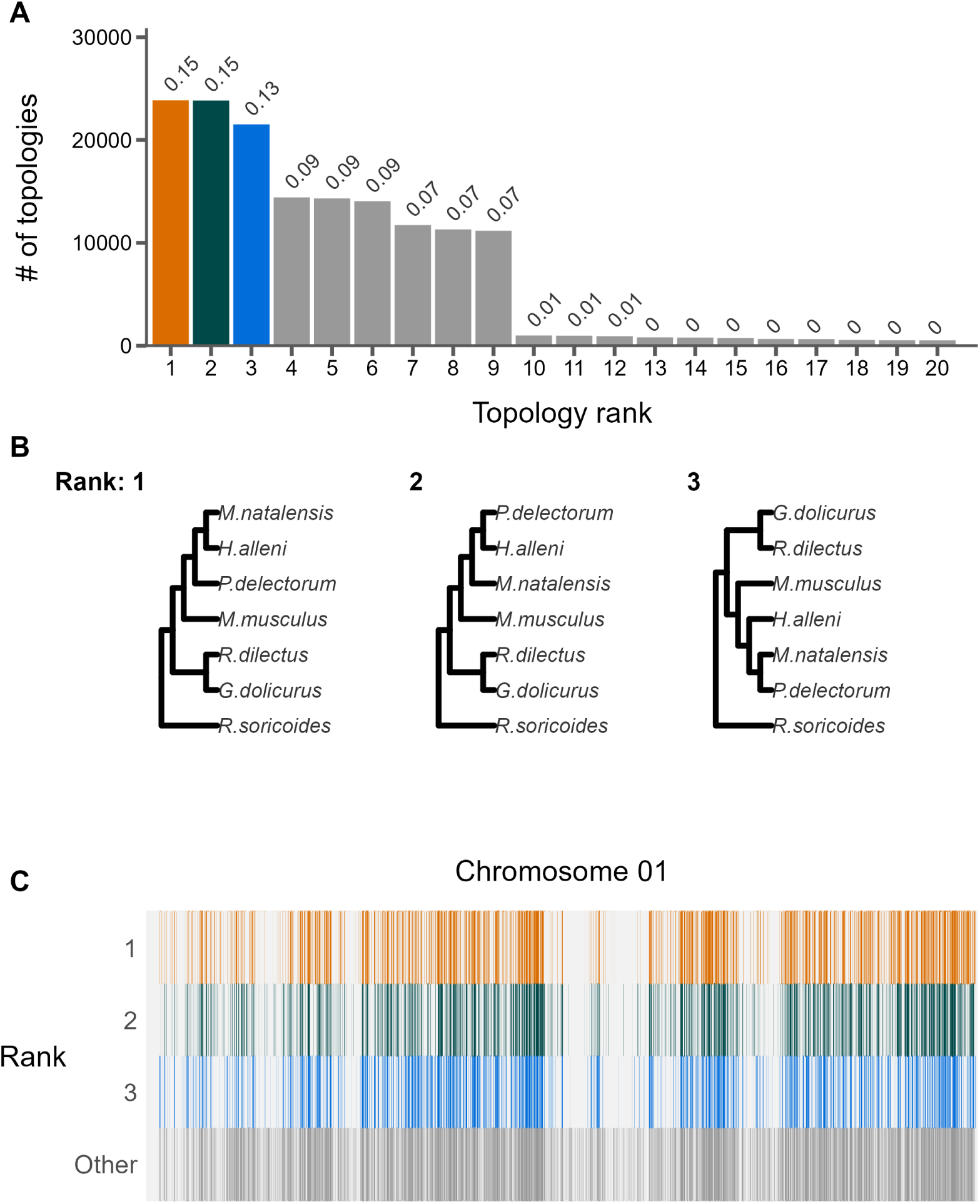
The landscape and profile of phylogenetic discordance across non-overlapping 10kb windows in murine genomes. A) Distribution of the 20 most frequent topologies recovered across all windows. Numbers above bars indicate proportion of each topology. B) The top three topologies recovered across all chromosomes 1. C) Distribution of the topologies recovered along chromosome 1. The x-axis is scaled to the length of the chromosome and each vertical bar represents one 10kb window. The three most frequent topologies occupy the first three rows while all other topologies are shown in the bottom row.

**Table 3:** The most frequently recovered topologies across all 10kb windows. RS = Rhyncomys soricoides, GD = Grammomys dolichurus, RD = Rhabdomys dilectus, MM = Mus musculus, HA = Hylomyscus alleni, MN = Mastomys natalensis, PD = Praomys delectorum.

| Rank | Topology | # of windows | Proportion of windows |
| --- | --- | --- | --- |
| 1 | (RS,((GD,RD),(MM,((HA,MN),PD)))); | 23864 | 0.146 |
| 2 | (RS,((GD,RD),(MM,((HA,PD),MN)))); | 23836 | 0.146 |
| 3* | (RS,((GD,RD),(MM,(HA,(MN,PD)))); | 21509 | 0.131 |
| 4 | (RS,(MM,((GD,RD),((HA,MN),PD)))); | 14417 | 0.088 |
| 5 | (RS,(MM,((GD,RD),((HA,PD),MN)))); | 14321 | 0.0874 |
| 6 | (RS,(MM,((GD,RD),(HA,(MN,PD)))); | 14044 | 0.0858 |
| 7 | (RS,(((HA,PD),MN),(MM,(GD,RD)))); | 11723 | 0.0716 |
| 8 | (RS,(((HA,MN),PD),(MM,(GD,RD)))); | 11308 | 0.0691 |

\*The topology recovered from concatenation of genes or UCEs

**Table 4:** Summaries of phylogenies per chromosome.

| Chromosome | # of unique topologies recovered |
| --- | --- |
| 1 | 184 |
| 2 | 123 |
| 3 | 114 |
| 4 | 144 |
| 5 | 134 |
| 6 | 172 |
| 7 | 218 |
| 8 | 133 |
| 9 | 116 |
| 10 | 110 |
| 11 | 93 |
| 12 | 179 |
| 13 | 186 |
| 14 | 173 |
| 15 | 96 |
| 16 | 94 |
| 17 | 188 |
| 18 | 75 |
| 19 | 82 |
| X | 207 |

While relationships among windows varied greatly and visual inspection revealed no clear partitioning of topological structures along the chromosome (*e.g.*, Fig. 2C), we found that phylogenies were not randomly distributed across chromosomes. We measured tree distance between adjacent windows using the weighted Robinson-Foulds metric and compared these distances to those measured between randomly selected windows on different chromosomes. We found that tree similarity between windows decayed logarithmically along chromosomes (Fig. 3A and B) and the distance at which tree similarity appeared random varied among chromosomes from 0.15 Megabases (Mb) on chromosome 17 to 141.29 Mb on the chromosome 2 (Fig. 3C, Fig. S5). While chromosomes 2, 7, 9, and 11 were autosomal outliers with distances between windows to random-like trees exceeding 25 Mb, the average distance among all other autosomes was only 2.1 Mb. The rates at which phylogenetic similarity decayed tended to be inversely proportional to the distance at which two randomly drawn phylogenies lost similarity (Fig. 3D).

**Figure 3:**
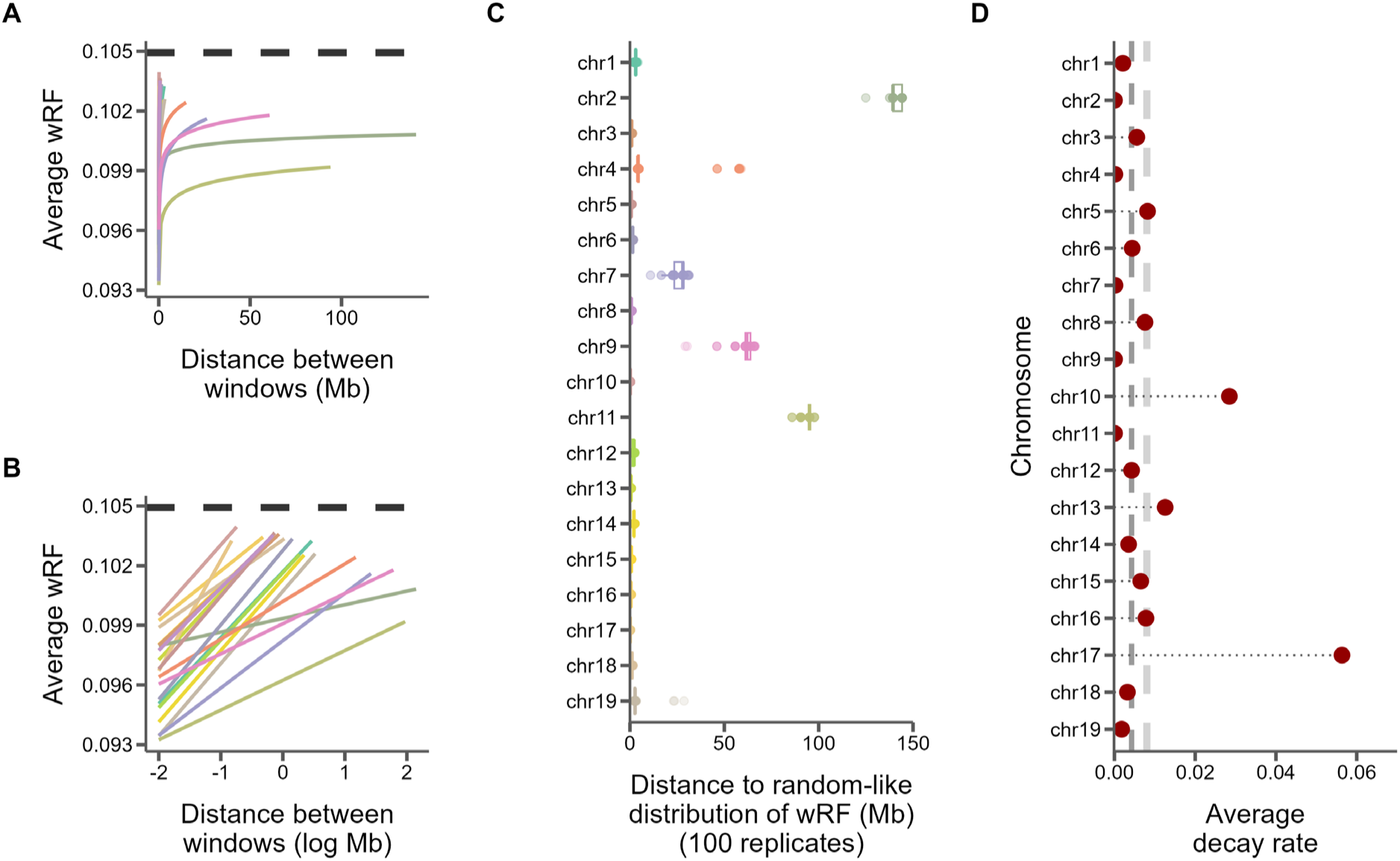
Similarity between 10kb windows decays as genomic distance between windows increases. A) The log fit to the mean of distributions of weighted Robinson-Foulds distances between trees of windows at increasing genomic distance (10kb steps). Each line represents one chromosome. B) The same, but on a log scale with a linear fit. C) For every window on each chromosome, the genomic distance between windows at which tree distance becomes random for 100 replicates of random window selection. D) The slopes of the correlation between genomic distance and tree distance from panel B represent the rate at which tree similarity decays across the genome.

To assess how large structural variation, such as inversions and translocations, may influence our inferences of phylogenetic relatedness along the genome, we aligned the reference genomes of mouse and rat. These two species span the divergence of the sample for which we assessed genome-wide discordance so the level of large structural variation present among them should give us an idea of the amount of ancestral variation in our sample. We find that the mouse and rat genomes are largely co-linear for large, aligned chunks, with large translocations and inversions on mouse chromosomes 5, 8, 10, and 13 (Fig. S6). We also observe large-scale inversions on chromosome 16. We find that, while co-linearity of most chromosomes is conserved between mouse and rat, the size of the 300,000 aligned chunks averages under 10 kb, with the average distance between aligned segments being between 2,380 bp on the mouse genome and 4927 bp on the rat chromosome (Fig. S7). This has two major consequences for our results: 1) this prevented us from transposing the coordinate system from mouse to rat with enough resolution to use genetic maps from rat and 2) this means that most other structural variation in our sample is likely small insertions of transposable elements (SINES which are about 150-500 bp in length and LINES which are about 4-7kb in length (Platt et al. 2018)) that should have a negligible effect on our discordance analyses since our window size is much larger and we excluded windows that were made up of mostly repeats.

### Discordance with recombination rate and other genomic features

Using markers from genetic crosses within *M. musculus* (Shifman et al. 2006; Cox et al. 2009) we examined whether regions of the genome with high recombination also showed more phylogenetic discordance over short genetic distances than regions with low recombination. Specifically, for each 5 Mb window we calculated recombination rate as the relationship between map location and the physical location for all markers within the window. Then within each 5 Mb window for which we had calculated recombination rate (Fig. S8), we measured tree similarity between the first and last 10 kb window. Surprisingly, we found no relationship between tree similarity and recombination rates measure at this scale (Fig. 4). We do, however, find that regions of the genome centered on recombination hotspots identified in *M. musculus* have a significantly slower rate of decay in similarity over genomic distance compared to windows that are not centered on hotspots (*p* = 0.019; Fig. 5A) and they are also significantly more phylogenetically similar over short distances (*p* = 0.015 Fig. 5B). This indicates that regions around hotspots retain a higher degree of phylogenetic similarity than other regions of the genome.

**Figure 4:**
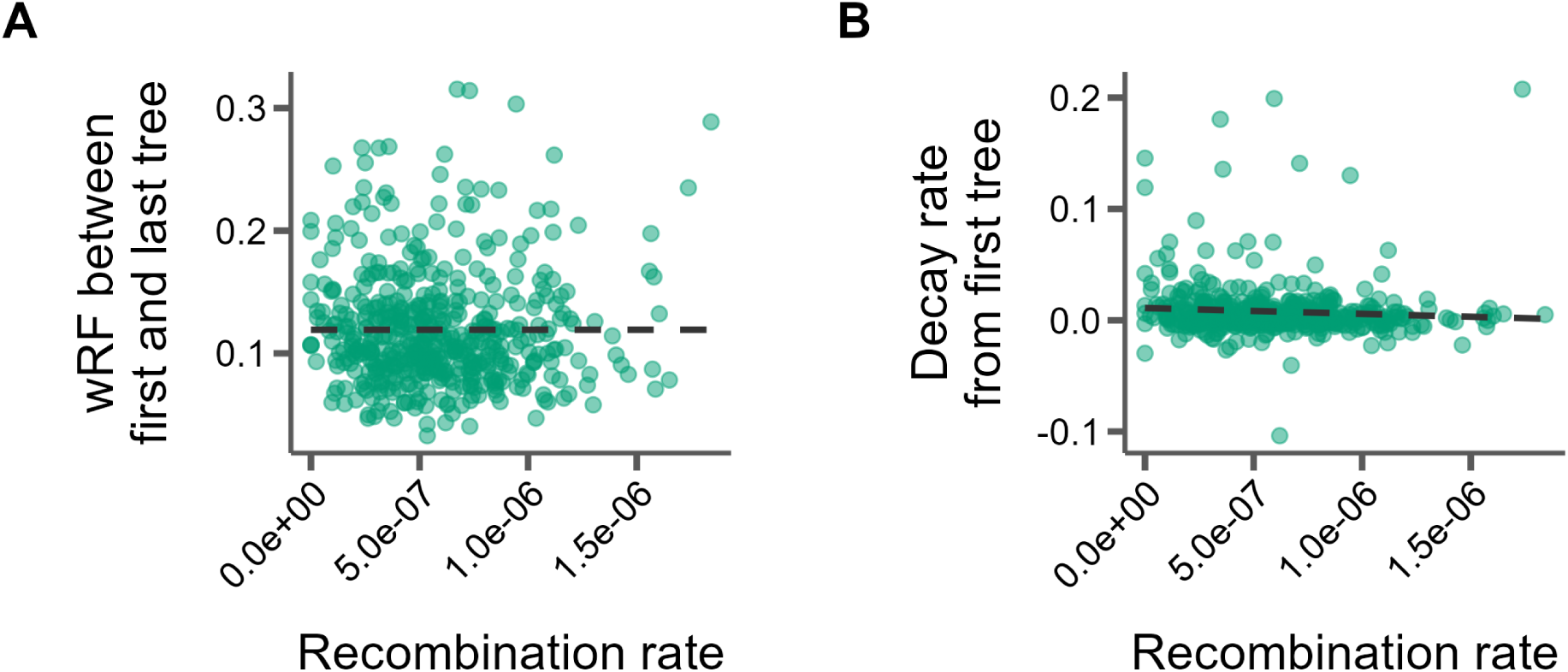
No correlation between tree similarity and recombination rate in 5Mb windows. A) Tree similarity as measured by the weighted Robinson-Foulds distance between the first and last 10kb windows within the 5Mb window. B) The slopes of the linear correlation between the weighted Robinson-Foulds distances between the first 10kb window and every other 10kb window within a 5Mb window represent the rate at which tree similarity decays over each 5Mb window.

**Figure 5:**
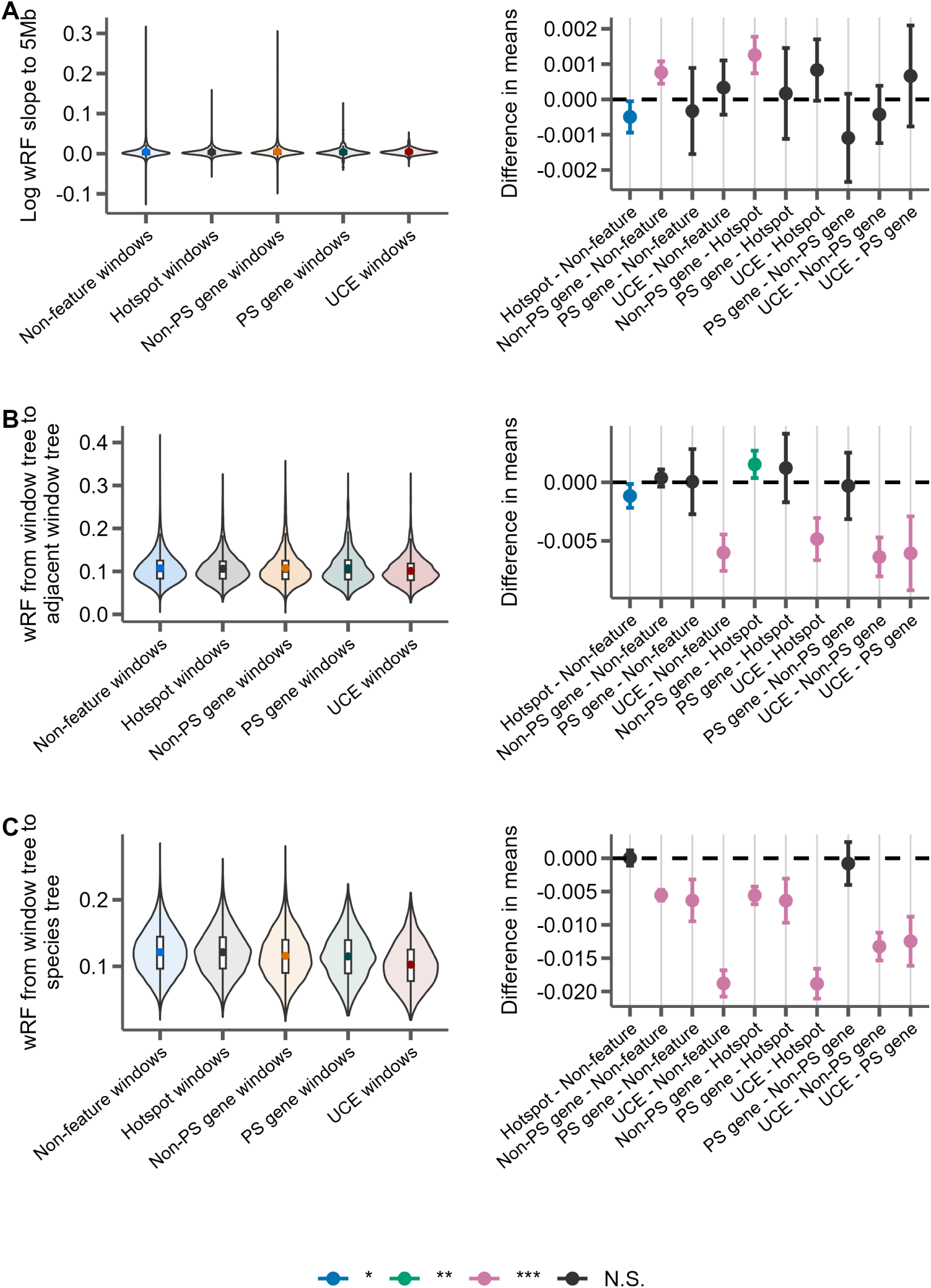
Distributions of weighted Robinson-Foulds distance from trees constructed from 10kb windows either centered on recombination hotspots (Hotspot), protein-coding genes without evidence for positive selection (Non-PS genes), protein coding genes with evidence for positive selection (PS genes), UCEs, or containing none of these features (Non-feature). For each panel, the left portion shows the distributions of the measure for each feature type and the right panel shows the differences in means for each pairwise comparison of features with significance assessed with Tukey’s range test. The labels on the x-axis indicate the feature pairs being compared, with the first feature being the reference (*i.e.* points above 0 indicate this feature has a higher mean). *P-*value thresholds: * < 0.05, ** < 0.01, *** <0.001. A) The rate of decay of phylogenetic similarity is calculated as the slope of a linear regression between wRF and the log distance between each window up to 5Mb away from the feature window. B) The phylogenetic similarity of windows immediately adjacent to feature windows. C) The phylogenetic similarity between the species tree inferred from protein-coding gene trees and the feature window.

Evolutionary relationships around certain conserved genomic features may also be shaped by locally reduced effective population sizes due to a history of pervasive linked negative or positive selection. To test for this, we measured tree similarity in 10 kb windows around all annotated protein coding genes, ultra-conserved elements (UCEs), and protein coding genes identified as evolving rapidly (*i.e.*, significantly elevated *dN*/*dS*) due to positive directional selection out to 5Mb and compared these patterns relative to chromosome-wide trends. We find that the phylogenetic similarity around protein coding genes is similar to that of windows without any genomic features (Fig. 5B), but that this similarity decays more rapidly around genes (*p* = 6.38e-8; Fig. 5A). Notably, we find the opposite to be true of UCEs, which have immediately adjacent regions that are much more similar than regions surrounding other features like hotspots (*p* = 2.42e-12), coding genes (*p* = 4.65e-14), and rapidly evolving coding genes (*p* = 1.56e-6), as well as windows that do not include any of these features (*p* = 5.02e-14; Fig. 5B) while decaying at roughly equivalent rates as these features with increasing genomic distance (Fig. 5A). In other words, regions around recombination hotspots have unexpectedly high phylogenetic similarity farther away from the hotspot while regions immediately surrounding UCE’s have unexpectedly high phylogenetic similarity both in the immediately adjacent regions and over long distances. We also find that the 10kb windows centered on most features differ in how similar they are to the species tree as inferred from coding genes or UCEs alone. All features except recombination hotspots are more similar to the species tree on average than windows that contain no features, while UCEs are more similar to the species tree than when compared to any other feature (Fig. 5C). We also note that positively selected genes are significantly more similar to the species tree than recombination hotspots, and genes, whether positively selected for or not, are equally similar to the species tree inferred from them (Fig. 5C).

### Consequences of tree misspecification on analyses of molecular evolution

Given that tree similarity decreases as a function of genomic distance, we next examined how this relates to the evolution of protein coding sequences. Among a set of 22,261 protein coding transcripts annotated from the *M. musculus* genome, we calculated that the average distance between the start and end of the coding sequence was 37.02 kb, or roughly 4 non-overlapping 10 kb windows. At this distance, tree similarity is predicted to diminish considerably (*e.g.*, by 0.10 wRF units), meaning that the phylogenetic history of individual genes may often contain phylogenetic discordance (Mendes and Hahn 2016). We also found that out of the 67,566 times the coding sequence in a gene overlapped with a 10 kb window, the inferred topology of the gene tree exactly matched the topology of the corresponding window tree only 11% of the time. Thus, the common practice of inferring gene trees on concatenated coding exons from a single transcript is still likely averaging over multiple possible histories.

With knowledge of how tree similarity varies across the genome, we tested how tree misspecification might impact standard *dN*/*dS* based phylogenetic analyses for positive directional selection. Specifically, we used the still common practice of assuming a single species tree for all genes and compared that to using individually inferred gene trees in three common statistical tests for positive selection: PAML’s M1a vs. M2a test (Yang 2007), HyPhy’s BUSTED test (Murrell et al. 2015), and HyPhy’s aBSREL test (Smith et al. 2015a). We found evidence that tree misspecification likely induces both false positive (type I) and false negative (type II) errors. For example, many genes were inferred as having experienced positive directional selection when using a single species tree, but not when using local gene trees and vice versa (Fig. 6A). Assuming the locally inferred gene tree is more accurate, this resulted in varying rates and types of error (Table 5). For BUSTED, we observe that 28% of genes inferred as having evolved under positive directional selection when using the gene tree were not inferred when using the concatenated species tree (likely false negatives). The opposite was true for M1a vs. M2a, where, among showing signals of positive selection in one of the two scenarios, 76% do so when using the concatenated species tree but not individual gene trees (likely false positives). In general, genes found to be evolving under positive selection using both tree types tended to be more concordant with the species tree than those that had evidence for positive selection either using only the concatenated tree or the gene tree (Fig. 6).

**Figure 6:**
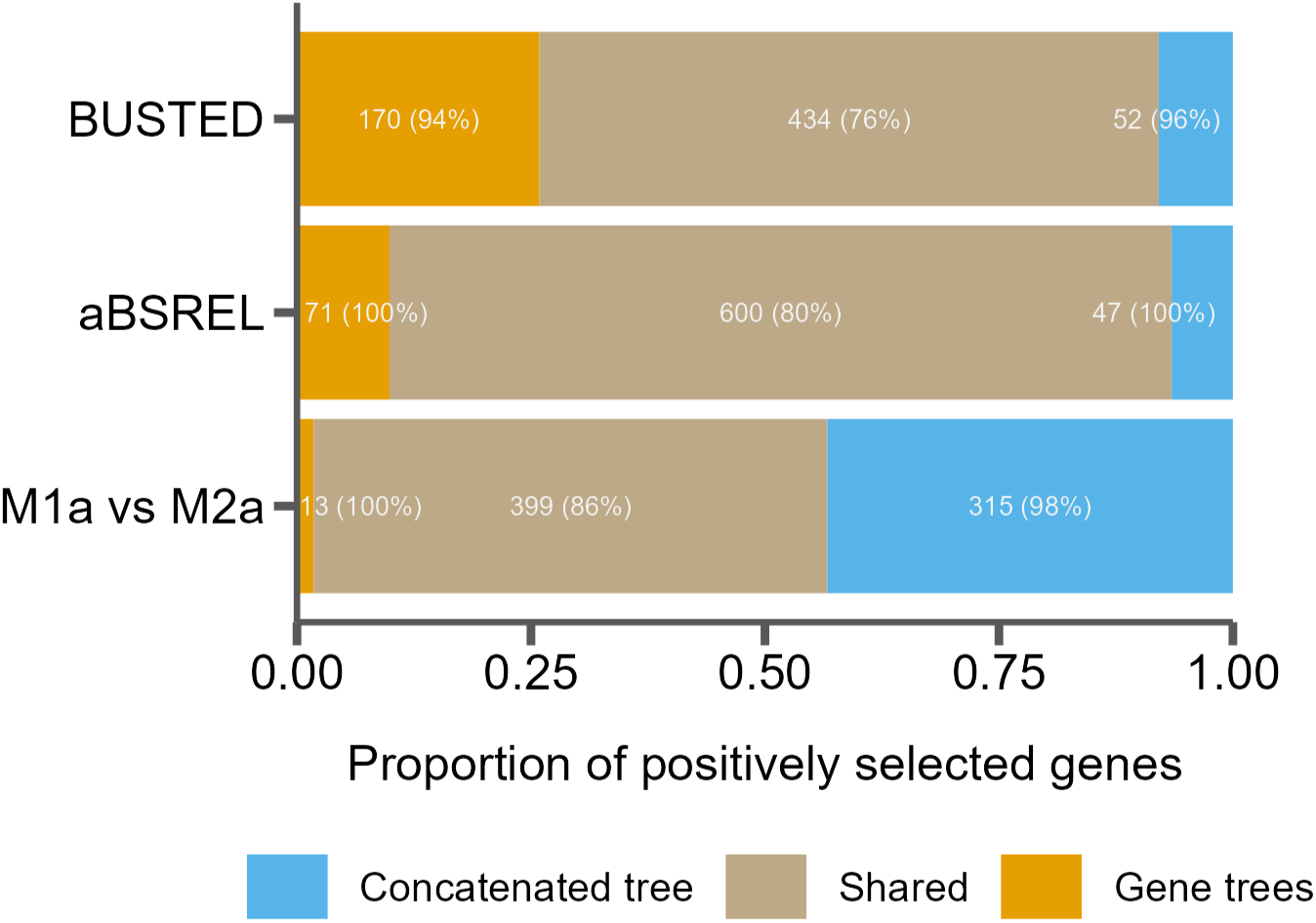
Tree misspecification leads to erroneous results in tests for positive selection. The proportion of genes inferred to be under positive selection for three tests using either a single species tree (concatenated tree) or individual gene trees, as well as those found in both cases (shared). Numbers in the bars indicate raw counts, and percentages indicate the percent of genes in that category that are discordant from the species tree.

**Table 5:** Rates and types of error when using concatenated trees for gene-based selection tests.

| Test | False positive rate | False negative rate |
| --- | --- | --- |
| BUSTED | 0.45% | 28.10% |
| aBSREL | 0.41% | 10.60% |
| M1a vs. M2a | 2.66% | 3.20% |

## Discussion

Phylogenies provide insight into the relationships of species and serve as a framework for asking questions about molecular and trait evolution. However, phylogenetic histories can vary extensively across regions of a genome, and evolutionary relationships between species may not often be well represented by a single representative species-level phylogeny. Here, we combine the resources of the house mouse (*Mus musculus*), a major model organism in biological and biomedical research with new genomes from seven closely related species to understand the systematics of murine rodents and causes and consequences of phylogenetic discordance along the murine genome. These new data and analyses help to place this important model system in an evolutionary context and begin to fill the gap in sampling of murine rodents which, despite their exceptional morphological and ecological diversity and species richness, have had relatively few whole genomes sequenced. They further provide us with the resources to study the landscape of phylogenetic discordance across the genome, understand how recombination and natural selection shape phylogenetic histories, and evaluate how assuming a single evolutionary history can compromise the study of molecular evolution. Beyond studying the patterns of discordance, this work highlights the importance of a nuanced molecular evolution analysis in a biomedical model system.

### Phylogenomic relationships of murine rodent lineages from conserved genomic regions

The extraordinary species richness of murine rodents complicates phylogenetic analyses because of the resources required to sample, sequence, and analyze such many widely distributed taxa. Earlier work either attempted to resolve specific groups such as *Mus* (Lundrigan et al. 2002; Suzuki et al. 2004) and *Apodemus* (Serizawa et al. 2000; Liu et al. 2004), or to uncover broader relationships across the subfamily (Martin et al. 2000; Steppan et al. 2005), in both cases using a limited number of nuclear or mitochondrial genes. Even so, evidence from these datasets suggested phylogenetic discordance across Murinae, including between mitochondrial and nuclear genes, that lead to different genomic regions supporting incompatible phylogenetic reconstructions (Suzuki et al. 2004; Steppan et al. 2005; White et al. 2009). Lecompte et al. (2008) provided one of the earliest well-supported phylogenetic reconstructions from across Murinae and the tribal classifications they proposed remain generally supported. More recent work has increased the number of taxa sampled, both for analyses of Murinae specifically (Pagès et al. 2016) and for their placement within Muridae and Muroidea (Schenk et al. 2013; Steppan and Schenk 2017; Rowe et al. 2019), but the number of loci used for phylogenetic inference remained limited to six loci or fewer. Recent work with a focus on Hydromyini made use of 1,245 (Roycroft et al. 2020) and 1,360 (Roycroft et al. 2021) exons for phylogenetic reconstruction.

We reconstructed a species tree of murine rodents based 2,632 UCEs from 18 species across the radiation. The inferred tree (Fig. 1) is topologically consistent with those inferred in previous studies (Lecompte et al. 2008; Steppan and Schenk 2017; Aghova et al. 2018). Branch support as estimated by UFBoot and SH-aLRT was uniformly high, and gene trees unambiguously support the tribal classification of Lecompte et al. (2008). However, four shorter branches show substantial gene tree discordance (Fig. 1, branches D, E, H, and J), with two recovered clades (E and J) being supported by less than half of all gene trees. Where phylogenetic reconstructions have used genomic markers – including UCEs – in other vertebrate taxa, gene tree discordance has been driven by a combination of incomplete lineage sorting or introgression (Alexander et al. 2017; Chan et al. 2020; Vanderpool et al. 2020; Alda et al. 2021), and methods to characterize these events are in active development (Hibbins and Hahn 2022). We also estimated divergence times on our inferred species tree using 4 fossil calibration points (Table 2), recovering times that are roughly consistent with the relatively young estimates found by (Steppan and Schenk 2017) (see Supplement).

### The genomic landscape of phylogenetic discordance

Limiting the number and nature of the loci used to resolve species relationships is often useful to get a clear picture of the species history across many taxa. However, such targeted approaches may fail to capture the degree and genomic landscape of discordance. Our results highlighted these limitations and the general relationships between phylogenetic patterns and functional attributes of the genome in several interesting ways. Using resources from the *M. musculus* model system, we found that the species tree inferred from genes or UCEs was not the most common topology recovered across the genome and did not match evolutionary relationships inferred for nearly 90% of the genome. This result is driven mainly by discordance among the three Praomyini species (Fig. 1) sampled for this study, with each alternate topology occurring at a frequency of roughly 14% (Fig. 2). The equal frequencies of this topology along the chromosome suggest that incomplete lineage sorting is the main driver of this discordance. However, it should be eye-opening that the topology robustly inferred from genes or UCEs was only the third most frequent topology among 10kb windows across the entire genome. This highlights the importance of accounting for phylogenetic discordance when studying aspects of genome evolution. While a single inferred species tree is of interest while studying the speciation and taxonomy of a group, individual locus trees must be used when making inferences about the evolution of particular regions or features of the genome.

While we observed no clear clustering of topological structures along most chromosomes (*e.g.*, Fig. 3C), we found that this enormous range of phylogenetic variation was not randomly distributed within chromosomes. Perhaps most surprisingly, we did not observe a relationship between recombination rates in mice (*M. musculus*) and the spatial scale of phylogenetic discordance. This negative result suggests that recombination rate evolves sufficiently quickly that contemporary estimates do not track variation over deeper evolutionary timescales, or at least at the 5 Mb scale we considered. Similar to findings in great apes (Hobolth et al. 2007), these results suggest that even high-resolution genetic resources from a single model species may be insufficient to help predict the landscape of discordance in a phylogenetic sample spanning 12 million years of evolution. This stands in contrast to the well-known negative relationship between population-level nucleotide variation and recombination rates in several mammals, including house mouse (Cai et al. 2009; Geraldes et al. 2011; Corbett-Detig et al. 2015; Kartje et al. 2020).

One important caveat is that our reference-based methods assume collinearity between *Mus* and the other lineages we are comparing (*i.e.*, no structural variation), at least at the scale that we are comparing variation. Structural variation, including substantial variation in chromosome numbers, are likely to be widespread in rodents (Stanyon et al. 1999; Yalcin et al. 2011; Romanenko et al. 2012; Keane et al. 2014) and has the potential to skew our results when comparing tree similarity between regions of the genome using multiple species – a structural variant may place a mapped region from one genome into a different genomic context in another. We currently lack chromosome scale assemblies for non-*Mus* and *Rattus* species and generating such assemblies may prove limiting given that most tissue resources for this group are derived from natural history collections that often lack high molecular weight DNA needed for most long-read DNA sequencing technologies. To assess the potential of syntenic variation to skew our results, we aligned the mouse (mm10) and rat (rnor6) genomes with minimap2 and find high degrees of chromosomal synteny and co-linearity between the two (Fig S11). While this does not preclude large rearrangements in other lineages in our sample, since these two genomes span the timescale of our sample for our genome-wide analysis of discordance it does mean that ancestrally most regions will be colinear. We do find, however, that most continuously aligned chunks between mouse and rat are small, being shorter than 10 kb on average and that there is on average only 2000-4000 bp between aligned segments, indicative of insertions and deletions of LINES and SINES preventing alignment (Fig. S7). This is also consistent with Ensembl’s alignment between the two species (release 107; (Cunningham et al. 2022)) and means that even though the genomes may be largely co-linear, we were unable to use rat in our analysis. Using our alignment between mouse and rat, we also find no significant correlation between structural variants and recombination rate (Fig. S9) consistent with recent results suggesting that most large structural changes in rodents occur during postmeiotic spermatogenesis and are independent of recombination machinery (Alvarez-Gonzalez et al. 2022).

Despite the lack of correlation between recombination rate and phylogenetic similarity, we did find varying degrees of phylogenetic similarity around recombination hotspots and other genomic features, such as genes and UCEs, presumably reflecting variation in the strength and frequency of linked selection over time. Natural selection reduces the effective population size (*Ne*) of genomic regions through the processes of genetic hitchhiking of variation linked to the fixation of positively selected mutations (*i.e*., selective sweeps; (Smith and Haigh 1974; Kaplan et al. 1989) and the purging of deleterious mutations (*i.e.*, background selection; (Charlesworth et al. 1993; Hudson and Kaplan 1995)). Thus, variation on parameters dependent on *Ne* – such as standing levels of nucleotide variation and patterns of incomplete lineage sorting – should be influenced by linkage to functional elements.

Our data suggest that background selection is important in structuring variation due to incomplete lineage sorting. Regions of the genome around recombination hotspots and UCEs were more similar than randomly chosen regions, though at different scales. Regions immediately adjacent to recombination hotspots are, on average, more similar than regions around no genomic features (Fig. 5B), and this similarity is retained over long distances (up to 5Mb; Fig. 5A). For UCEs, the regions immediately adjacent to them are significantly more similar than regions around no genomic features and this similarity dissipated at rates similar to chromosome-wide levels meaning that even at distances of up to 5Mb, the area around these regions remained more similar at long distances. Notably, this pattern is true for regions around UCEs compared to any other genomic feature we studied. These results suggest that a history of strong purifying selection at UCEs (Katzman et al. 2007) more generally strongly skews patterns of discordance consistent with a persistent local reduction in *Ne*. One practical consequence of this is that phylogenetic inferences based on UCE markers would seem less prone to discordance and may provide cleaner estimates of species tree history than randomly chosen regions. This was indeed the case as windows centered on UCEs have a higher degree of similarity to the species tree than other genomic features and recover the species tree topology 17% of the time, compared to 13% genome-wide or 15% for protein coding genes. Also interesting to note is that even though we recovered the same species tree topology for the seven species used in the genomic window analysis with both UCEs and protein-coding gene trees (Fig. 1), windows centered on UCEs are much more similar to the species tree than windows centered on protein coding genes (Fig. 5C). However, UCEs will also provide a skewed and potentially misleading view of levels of genome-wide discordance. Given this relationship, inferences based on UCEs may not, for example, be extended to related phylogenetic parameters of interest (e.g., ancestral population sizes), and, despite the relative ease of generating UCE data, such markers are not suitable for genetic inferences within populations.

The direction and frequency of selection also likely matters when considering the impact on variation in phylogenetic signal. Our focus on models of molecular evolution also allowed us to contrast patterns at genes experiencing positive directional selection to UCEs and genes evolving predominantly under purifying selection. Positive directional selection is generally assumed to be rare and punctuated across lineages (Kimura 1968; Ohta 1973; Fay 2011) while strong background selection at UCEs is likely to be more pervasive (Chen et al. 2007; Katzman et al. 2007), likely impacting the predictability of the influence of both forms of selection on discordance over evolutionary timescales. Consistent with this, we observed less conservation of phylogenetic signal on average surrounding rapidly evolving genes and more variance. The models that we considered are designed to detect recurrent positive directional selection. Nonetheless, the reduced diversity associated with these recurrent selective sweeps of beneficial amino acid substitutions at these genes appears to fade relatively quickly over time, at least relative to more pervasive purifying selection at UCEs and other gene regions.

### Discordance and Molecular Evolution

We also found that the choice of tree topology drastically affects the results from various common tests for positive selection. It is known that tree misspecification can lead to incorrect placement of substitutions on branches, possibly leading to spurious results for tests of positive directional selection (Hahn and Nakhleh 2016; Mendes and Hahn 2016). Here, we show that these errors result in false positive (detected signal for selection only when using the gene tree) and false negative results (detected signal for selection only when using the species tree).

For each of three selection tests run, HyPhy’s BUSTED and aBSREL and PAML’s M1a vs. M2asome genes showed evidence of positive selection whether the species tree or gene tree was used while many other genes had signatures of positive selection restricted only to a single tree. The genes unique to the type of tree used were often discordant with the species tree. In contrast, the genes that showed evidence of positive selection regardless of tree had levels of discordance comparable to all genes (85%, Figure 6, numbers in parentheses). This suggests that mis-mapping substitutions by supplying these tests with the wrong tree (*i.e.*, the concatenated species tree when gene trees are discordant) can lead to inflated false positive and false negative rates when inferring genes under positive selection. The magnitude and direction of these biases were dependent on the underlying model. So-called branch-site models, such as HyPhy’s BUSTED and aBSREL models, that allow substitution rates to vary among both branches and codon sites, resulted in more genes inferred with evidence for positive selection when using the inferred gene (*i.e.*, the correct tree, assuming no errors in gene tree reconstruction). This means that using a concatenated tree for these tests reduces the power to detect positive selection. On the other hand, models that only allow rates to vary among sites, such as PAML’s M1a vs. M2a test, showed an increase in the number of false positives inferred when using the wrong tree. That is, tree mis-specification results in spurious increases in *dN/dS* that mimic positive selection.

These results likely have wide-ranging implications for phylogenetics and comparative genomic analysis. First, it is imperative that when testing a specific locus for positive selection, discordance among loci must be accounted for. This is most easily achieved by simply using the gene tree (or other locus type) as input to the test for selection (Mendes and Hahn 2016; Roycroft et al. 2021). However, as Mendes and Hahn (2016) pointed out, this may not completely mask the effects of discordance on substitution rates, as sites within a single gene may still have evolved under different histories because of within-gene recombination. We also find evidence for this here, given that tree similarity diminished at scales that are less than the average genomic distance between the beginning and end of a coding sequence in mice (37.02 kb in this data set). Nevertheless, starting with an inferred gene tree is advisable whenever possible, followed by a secondary analysis of evidence for within-gene variation in phylogenetic history. However, incorporating discordance into a comparative framework is not trivial and many comparative genomic methods assume a single species tree that test for changes in substitution rates in a phylogeny (Pollard et al. 2010; Hu et al. 2019; Partha et al. 2019). Even methods that allow the use of different trees for different loci (like PAML and HyPhy) are still commonly applied with a single species tree across loci (Carbone et al. 2014; Foote et al. 2015; van der Valk et al. 2021; Treaster et al. 2023). Our results confirm that the use of a single tree for such tests that rely on accurate estimation of substitution rates are likely to lead to both false negative and false positive inferences of positive selection.

Finally, because of recombination’s underlying contribution to phylogenomic discordance, one might be tempted to control for alternate topologies in a comparative genomic dataset by using a genetic map from a single, well-studied species and sampling regions with low recombination rates. However, our results show that using a single recombination map is insufficient to control for discordance even among a small sample of seven species. This is likely because the recombination rate also evolves and is linked to structural variation (Morgan et al. 2017) which is unaccounted for with use of a single reference genome, resulting in a diminished correlation signal between rates and phylogenetic discordance over time. This would likely be compounded for more species spanning deeper evolutionary timescales.

## Conclusions

Overall, our results increase whole genome sampling in the diverse group of murine rodents and provide a broad murine species tree based on thousands of UCEs. The high-quality *M. musculus* genome allows us to examine fine-scale patterns and effects of phylogenetic discordance along chromosomes, revealing how discordance at evolutionary timescales varies with genome biology. We also demonstrate how phylogenetic discordance can mislead common tests for selection if only a single species tree is used. Using our results, we can better understand the complexities of phylogenomic datasets which will help to ensure that steps are taken to accommodate these details in future comparative studies.

## Acknowledgments

We thank Jake Esselstyn and Kevin Rowe for helpful comments and discussion on the murine species tree. We also thank Brant Faircloth and Trevor Sless for advice on using phyluce. We are grateful for tissue samples provided by Chris Conroy at the Museum of Vertebrate Zoology, Berkeley, CA (MVZ) and Adam Ferguson at the Field Museum of Natural History, Chicago, IL (FMNH), and to the original collectors. This work was supported by the National Science Foundation (DEB-1754096 to J.M.G), the Eunice Kennedy Shriver National Institute of Child Health and Human Development of the National Institutes of Health (R01-HD094787 to J.M.G.). J.J.H. received financial support from the Cornell Center for Vertebrate Genomics. J. S. B. was supported by the University of Michigan Life Sciences Fellows program and the Jean Wright Cohn Endowment Fund at the University of Michigan Museum of Zoology. Computations for species tree reconstruction were performed using the computer clusters and data storage resources of the University of California Riverside HPCC, which were funded by grants from NSF (MRI-2215705, MRI-1429826) and NIH (1S10OD016290-01A1), and the Cornell University Biotechnology Resource Center BioHPC (RRID:SCR_021757) with help from Qi Sun. Bioinformatic analyses for genomic discordance and selection tests were conducted using the University of Montana Griz Shared Computing Cluster supported by grants from the NSF (CC-2018112 and OAC-1925267, J.M.G. co-PI). Any opinions, findings, and conclusions or recommendations expressed in this material are those of the authors and do not necessarily reflect the views of the NSF or the NIH.

## Data availability

All raw reads and assembled genomes are available on XX. Pseudo-assemblies and locus alignments (UCEs, genes, and windows) are available on XX. All code and summary data for this project are deposited in https://github.com/gwct/murine-discordance.

## References

Pdb (the paleobiology database) [Internet]. 2011 January 21st, 2022]. Available from: http://paleodb.org/

Aghova T, Kimura Y, Bryja J, Dobigny G, Granjon L, Kergoat GJ. 2018. Fossils know it best: Using a new set of fossil calibrations to improve the temporal phylogenetic framework of murid rodents (rodentia: Muridae). Mol Phylogenet Evol. 128:98–111.

Alda F, Ludt WB, Elias DJ, McMahan CD, Chakrabarty P. 2021. Comparing ultraconserved elements and exons for phylogenomic analyses of middle american cichlids: When data agree to disagree. Genome Biol Evol. 13.

Alexander AM, Su YC, Oliveros CH, Olson KV, Travers SL, Brown RM. 2017. Genomic data reveals potential for hybridization, introgression, and incomplete lineage sorting to confound phylogenetic relationships in an adaptive radiation of narrow-mouth frogs. Evolution. 71:475–488.

Alvarez-Gonzalez L, Burden F, Doddamani D, Malinverni R, Leach E, Marin-Garcia C, Marin-Gual L, Gubern A, Vara C, Paytuvi-Gallart A, et al. 2022. 3d chromatin remodelling in the germ line modulates genome evolutionary plasticity. Nat Commun. 13:2608.

Avise JC, Robinson TJ. 2008. Hemiplasy: A new term in the lexicon of phylogenetics. Syst Biol. 57:503–507.

Baum DA. 2007. Concordance trees, concordance factors, and the exploration of reticulate genealogy. TAXON. 56:417–426.

Benjamini Y, Hochberg Y. 1995. Controlling the false discovery rate: A practical and powerful approach to multiple testing. Journal of the Royal statistical society: series B (Methodological). 57:289–300.

Bloom BH. 1970. Space/time trade-offs in hash coding with allowable errors. *Commun. ACM*. 13:422–426. Böcker S, Canzar S, Gunnar WK. 2013. The generalized robinson-foulds metric. In: Darling A, Stoye J, editors. Algorithms in bioinformatics. Berlin, Heidelberg: Springer.

Bolger AM, Lohse M, Usadel B. 2014. Trimmomatic: A flexible trimmer for illumina sequence data. Bioinformatics. 30:2114–2120.

Borowiec ML. 2016. Amas: A fast tool for alignment manipulation and computing of summary statistics. PeerJ. 4:e1660.

Bradley RK, Roberts A, Smoot M, Juvekar S, Do J, Dewey C, Holmes I, Pachter L. 2009. Fast statistical alignment. PLoS Comput Biol. 5:e1000392.

Cai JJ, Macpherson JM, Sella G, Petrov DA. 2009. Pervasive hitchhiking at coding and regulatory sites in humans. PLoS Genet. 5:e1000336.

Capella-Gutierrez S, Silla-Martinez JM, Gabaldon T. 2009. Trimal: A tool for automated alignment trimming in large-scale phylogenetic analyses. Bioinformatics. 25:1972–1973.

Carbone L, Harris RA, Gnerre S, Veeramah KR, Lorente-Galdos B, Huddleston J, Meyer TJ, Herrero J, Roos C, Aken B, et al. 2014. Gibbon genome and the fast karyotype evolution of small apes. Nature. 513:195–201.

Chan KO, Hutter CR, Wood PL, Jr., Grismer LL, Brown RM. 2020. Target-capture phylogenomics provide insights on gene and species tree discordances in old world treefrogs (anura: Rhacophoridae). Proc Biol Sci. 287:20202102.

Charlesworth B, Morgan MT, Charlesworth D. 1993. The effect of deleterious mutations on neutral molecular variation. Genetics. 134:1289–1303.

Chen CT, Wang JC, Cohen BA. 2007. The strength of selection on ultraconserved elements in the human genome. Am J Hum Genet. 80:692–704.

Chernomor O, von Haeseler A, Minh BQ. 2016. Terrace aware data structure for phylogenomic inference from supermatrices. Syst Biol. 65:997–1008.

Corbett-Detig RB, Hartl DL, Sackton TB. 2015. Natural selection constrains neutral diversity across a wide range of species. PLoS Biol. 13:e1002112.

Cox A, Ackert-Bicknell CL, Dumont BL, Ding Y, Bell JT, Brockmann GA, Wergedal JE, Bult C, Paigen B, Flint J, et al. 2009. A new standard genetic map for the laboratory mouse. Genetics. 182:1335–1344.

Cunningham F, Allen JE, Allen J, Alvarez-Jarreta J, Amode MR, Armean IM, Austine-Orimoloye O, Azov AG, Barnes I, Bennett R, et al. 2022. Ensembl 2022. Nucleic Acids Res. 50:D988–D995.

Danecek P, Bonfield JK, Liddle J, Marshall J, Ohan V, Pollard MO, Whitwham A, Keane T, McCarthy SA, Davies RM, et al. 2021. Twelve years of samtools and bcftools. Gigascience. 10.

Degnan JH, Rosenberg NA. 2006. Discordance of species trees with their most likely gene trees. PLoS Genet. 2:e68.

Edwards SV. 2009. Is a new and general theory of molecular systematics emerging? Evolution. 63:1–19.

Faircloth BC. 2013. Illumiprocessor: A trimmomatic wrapper for parallel adapter and quality trimming.

Faircloth BC. 2016. Phyluce is a software package for the analysis of conserved genomic loci.Bioinformatics. 32:786–788.

Faircloth BC, McCormack JE, Crawford NG, Harvey MG, Brumfield RT, Glenn TC. 2012. Ultraconserved elements anchor thousands of genetic markers spanning multiple evolutionary timescales. Syst Biol. 61:717–726.

Fay JC. 2011. Weighing the evidence for adaptation at the molecular level. Trends Genet. 27:343–349.

Fontaine MC, Pease JB, Steele A, Waterhouse RM, Neafsey DE, Sharakhov IV, Jiang X, Hall AB, Catteruccia F, Kakani E, et al. 2015. Mosquito genomics. Extensive introgression in a malaria vector species complex revealed by phylogenomics. Science. 347:1258524.

Foote AD, Liu Y, Thomas GW, Vinar T, Alfoldi J, Deng J, Dugan S, van Elk CE, Hunter ME, Joshi V, et al. 2015. Convergent evolution of the genomes of marine mammals. Nat Genet. 47:272–275.

Geraldes A, Basset P, Smith KL, Nachman MW. 2011. Higher differentiation among subspecies of the house mouse (mus musculus) in genomic regions with low recombination. Mol Ecol. 20:4722–4736.

Gibbs RA, Weinstock GM, Metzker ML, Muzny DM, Sodergren EJ, Scherer S, Scott G, Steffen D, Worley KC, Burch PE, et al. 2004. Genome sequence of the brown norway rat yields insights into mammalian evolution. Nature. 428:493–521.

Green RE, Krause J, Briggs AW, Maricic T, Stenzel U, Kircher M, Patterson N, Li H, Zhai W, Fritz MH, et al. 2010. A draft sequence of the neandertal genome. Science. 328:710–722.

Guindon S, Dufayard JF, Lefort V, Anisimova M, Hordijk W, Gascuel O. 2010. New algorithms and methods to estimate maximum-likelihood phylogenies: Assessing the performance of phyml 3.0. Syst Biol. 59:307–321.

Hahn MW, Nakhleh L. 2016. Irrational exuberance for resolved species trees. Evolution. 70:7–17.

Hibbins MS, Hahn MW. 2022. Phylogenomic approaches to detecting and characterizing introgression. Genetics. 220.

Hinrichs AS, Karolchik D, Baertsch R, Barber GP, Bejerano G, Clawson H, Diekhans M, Furey TS, Harte RA, Hsu F, et al. 2006. The ucsc genome browser database: Update 2006. Nucleic Acids Res. 34:D590–598.

Hoang DT, Chernomor O, von Haeseler A, Minh BQ, Vinh LS. 2018. Ufboot2: Improving the ultrafast bootstrap approximation. Mol Biol Evol. 35:518–522.

Hobolth A, Christensen OF, Mailund T, Schierup MH. 2007. Genomic relationships and speciation times of human, chimpanzee, and gorilla inferred from a coalescent hidden markov model. PLoS Genet. 3:e7.

Holm S. 1979. A simple sequentially rejective multiple test procedure. Scandinavian journal of statistics.65–70.

Hu Z, Sackton TB, Edwards SV, Liu JS. 2019. Bayesian detection of convergent rate changes of conserved noncoding elements on phylogenetic trees. Mol Biol Evol. 36:1086–1100.

Hudson RR. 1983. Testing the constant-rate neutral allele model with protein sequence data. Evolution. 37:203–217.

Hudson RR, Kaplan NL. 1988. The coalescent process in models with selection and recombination. Genetics. 120:831–840.

Hudson RR, Kaplan NL. 1995. Deleterious background selection with recombination. Genetics. 141:1605–1617.

Huson DH, Klöpper T, Lockhart PJ, Steel MA. 2005. Reconstruction of reticulate networks from gene trees. In: Miyano S, Mesirov J, Kasif S, Istrail S, Pevzner PA, Waterman M, editors. Research in computational molecular biology. RECOMB 2005. Lecture Notes in Computer Science: Springer, Berlin, Heidelberg.

Jackman SD, Vandervalk BP, Mohamadi H, Chu J, Yeo S, Hammond SA, Jahesh G, Khan H, Coombe L, Warren RL, et al. 2017. Abyss 2.0: Resource-efficient assembly of large genomes using a bloom filter. Genome Res. 27:768–777.

Jarvis ED, Mirarab S, Aberer AJ, Li B, Houde P, Li C, Ho SY, Faircloth BC, Nabholz B, Howard JT, et al. 2014. Whole-genome analyses resolve early branches in the tree of life of modern birds. Science. 346:1320–1331.

Jones MR, Mills LS, Alves PC, Callahan CM, Alves JM, Lafferty DJR, Jiggins FM, Jensen JD, Melo-Ferreira J, Good JM. 2018. Adaptive introgression underlies polymorphic seasonal camouflage in snowshoe hares. Science. 360:1355–1358.

Junier T, Zdobnov EM. 2010. The newick utilities: High-throughput phylogenetic tree processing in the unix shell. Bioinformatics. 26:1669–1670.

Kalyaanamoorthy S, Minh BQ, Wong TKF, von Haeseler A, Jermiin LS. 2017. Modelfinder: Fast model selection for accurate phylogenetic estimates. Nat Methods. 14:587–589.

Kaplan NL, Hudson RR, Langley CH. 1989. The "hitchhiking effect" revisited. Genetics. 123:887–899.

Kartje ME, Jing P, Payseur BA. 2020. Weak correlation between nucleotide variation and recombination rate across the house mouse genome. Genome Biol Evol. 12:293–299.

Katoh K, Standley DM. 2013. Mafft multiple sequence alignment software version 7: Improvements in performance and usability. Mol Biol Evol. 30:772–780.

Katzman S, Kern AD, Bejerano G, Fewell G, Fulton L, Wilson RK, Salama SR, Haussler D. 2007. Human genome ultraconserved elements are ultraselected. Science. 317:915.

Keane TM, Wong K, Adams DJ, Flint J, Reymond A, Yalcin B. 2014. Identification of structural variation in mouse genomes. Front Genet. 5:192.

Kimura M. 1968. Evolutionary rate at the molecular level. Nature. 217:624–626.

Kowalczyk A, Meyer WK, Partha R, Mao W, Clark NL, Chikina M. 2019. Rerconverge: An r package for associating evolutionary rates with convergent traits. Bioinformatics. 35:4815–4817.

Kulathinal RJ, Stevison LS, Noor MA. 2009. The genomics of speciation in drosophila: Diversity, divergence, and introgression estimated using low-coverage genome sequencing. PLoS Genet. 5:e1000550.

Kumon T, Ma J, Akins RB, Stefanik D, Nordgren CE, Kim J, Levine MT, Lampson MA. 2021. Parallel pathways for recruiting effector proteins determine centromere drive and suppression. Cell. 184:4904–4918 e4911.

Calculating and interpreting gene- and site-concordance factors in phylogenomics [Internet]. The Lanfear Lab @ ANU2018 September 20, 2021]. Available from: http://www.robertlanfear.com/blog/files/concordance_factors.html

Lecompte E, Aplin K, Denys C, Catzeflis F, Chades M, Chevret P. 2008. Phylogeny and biogeography of african murinae based on mitochondrial and nuclear gene sequences, with a new tribal classification of the subfamily. BMC Evol Biol. 8:199.

Lewontin RC, Birch LC. 1966. Hybridization as a source of variation for adaptation to new environments. Evolution. 20:315–336.

Li H. 2013. Aligning sequence reads, clone sequences and assembly contigs with bwa-mem. arXiv preprint arXiv:1303.3997.

Li H. 2018. Minimap2: Pairwise alignment for nucleotide sequences. Bioinformatics. 34:3094–3100.

Liu X, Wei F, Li M, Jiang X, Feng Z, Hu J. 2004. Molecular phylogeny and taxonomy of wood mice (genus apodemus kaup, 1829) based on complete mtdna cytochrome b sequences, with emphasis on chinese species. Mol Phylogenet Evol. 33:1–15.

Lopes F, Oliveira LR, Kessler A, Beux Y, Crespo E, Cardenas-Alayza S, Majluf P, Sepulveda M, Brownell RL, Franco-Trecu V, et al. 2021. Phylogenomic discordance in the eared seals is best explained by incomplete lineage sorting following explosive radiation in the southern hemisphere. Syst Biol. 70:786–802.

Lundrigan BL, Jansa SA, Tucker PK. 2002. Phylogenetic relationships in the genus mus, based on paternally, maternally, and biparentally inherited characters. Syst Biol. 51:410–431.

Maddison WP. 1997. Gene trees in species trees. Systematic Biology. 46:523–536.

Marks P, Garcia S, Barrio AM, Belhocine K, Bernate J, Bharadwaj R, Bjornson K, Catalanotti C, Delaney J, Fehr A, et al. 2019. Resolving the full spectrum of human genome variation using linked-reads. Genome Res. 29:635–645.

Martin Y, Gerlach G, Schlotterer C, Meyer A. 2000. Molecular phylogeny of european muroid rodents based on complete cytochrome b sequences. Mol Phylogenet Evol. 16:37–47.

McKenzie PF, Eaton DAR. 2020. The multispecies coalescent in space and time. bioRxiv.2020.2008.2002.233395.

Mendes FK, Fuentes-Gonzalez JA, Schraiber JG, Hahn MW. 2018. A multispecies coalescent model for quantitative traits. Elife. 7.

Mendes FK, Hahn MW. 2016. Gene tree discordance causes apparent substitution rate variation. Syst Biol. 65:711–721.

Mendes FK, Hahn Y, Hahn MW. 2016. Gene tree discordance can generate patterns of diminishing convergence over time. Mol Biol Evol. 33:3299–3307.

Minh BQ, Hahn MW, Lanfear R. 2020a. New methods to calculate concordance factors for phylogenomic datasets. Mol Biol Evol. 37:2727–2733.

Minh BQ, Schmidt HA, Chernomor O, Schrempf D, Woodhams MD, von Haeseler A, Lanfear R. 2020b. Iq-tree 2: New models and efficient methods for phylogenetic inference in the genomic era. Mol Biol Evol. 37:1530–1534.

Mölder F, Jablonski KP, Letcher B, Hall MB, Tomkins-Tinch CH, Sochat V, Forster J, Lee S, Twardziok SO, Kanitz A, et al. 2021. Sustainable data analysis with snakemake. F1000Res. 10:33.

Moore EC, Thomas GWC, Mortimer S, Kopania EEK, Hunnicutt KE, Clare-Salzler ZJ, Larson EL, Good JM. 2022. The evolution of widespread recombination suppression on the dwarf hamster (phodopus) x chromosome. Genome Biol Evol. 14.

Morgan AP, Gatti DM, Najarian ML, Keane TM, Galante RJ, Pack AI, Mott R, Churchill GA, de Villena FP. 2017. Structural variation shapes the landscape of recombination in mouse. Genetics. 206:603–619.

Mouse Genome Sequencing C, Waterston RH, Lindblad-Toh K, Birney E, Rogers J, Abril JF, Agarwal P, Agarwala R, Ainscough R, Alexandersson M, et al. 2002. Initial sequencing and comparative analysis of the mouse genome. Nature. 420:520–562.

Murrell B, Weaver S, Smith MD, Wertheim JO, Murrell S, Aylward A, Eren K, Pollner T, Martin DP, Smith DM, et al. 2015. Gene-wide identification of episodic selection. Mol Biol Evol. 32:1365–1371.

Nilsson P, Solbakken MH, Schmid BV, Orr RJS, Lv R, Cui Y, Song Y, Zhang Y, Baalsrud HT, Torresen OK, et al. 2020. The genome of the great gerbil reveals species-specific duplication of an mhcii gene. Genome Biol Evol. 12:3832–3849.

Ohta T. 1973. Slightly deleterious mutant substitutions in evolution. Nature. 246:96–98.

Pagès M, Fabre P-H, Chaval Y, Mortelliti A, Nicolas V, Wells K, Michaux JR, Lazzari V. 2016. Molecular phylogeny of south-east asian arboreal murine rodents. Zoologica Scripta. 45:349–364.

Pamilo P, Nei M. 1988. Relationships between gene trees and species trees. Mol Biol Evol. 5:568–583.

Paradis E, Schliep K. 2019. Ape 5.0: An environment for modern phylogenetics and evolutionary analyses in r. Bioinformatics. 35:526–528.

Partha R, Kowalczyk A, Clark NL, Chikina M. 2019. Robust method for detecting convergent shifts in evolutionary rates. Mol Biol Evol. 36:1817–1830.

Pease JB, Haak DC, Hahn MW, Moyle LC. 2016. Phylogenomics reveals three sources of adaptive variation during a rapid radiation. PLoS Biol. 14:e1002379.

Platt RN, 2nd, Vandewege MW, Ray DA. 2018. Mammalian transposable elements and their impacts on genome evolution. Chromosome Res. 26:25–43.

Pollard KS, Hubisz MJ, Rosenbloom KR, Siepel A. 2010. Detection of nonneutral substitution rates on mammalian phylogenies. Genome Res. 20:110–121.

Pond SL, Frost SD, Muse SV. 2005. Hyphy: Hypothesis testing using phylogenies. Bioinformatics. 21:676–679.

Poplin R, Ruano-Rubio V, DePristo MA, Fennell TJ, Carneiro MO, Auwera GAVd, Kling DE, Gauthier LD, Levy-Moonshine A, Roazen D, et al. 2018. Scaling accurate genetic variant discovery to tens of thousands of samples. bioRxiv.201178.

Quinlan AR, Hall IM. 2010. Bedtools: A flexible suite of utilities for comparing genomic features. Bioinformatics. 26:841–842.

R Core Team. 2021.R: A language and environment for statistical computing. Vienna, Austria.

Ranwez V, Douzery EJP, Cambon C, Chantret N, Delsuc F. 2018. Macse v2: Toolkit for the alignment of coding sequences accounting for frameshifts and stop codons. Mol Biol Evol. 35:2582–2584.

Revell LJ. 2012. Phytools: An r package for phylogenetic comparative biology (and other things). Methods in Ecology and Evolution. 3:217–223.

Robinson DF, Foulds LR. 1981. Comparison of phylogenetic trees. Mathematical Biosciences. 53:131–147.

Robinson DF, Foulds LR editors.; 1979 Berlin, Heidelberg.

Romanenko SA, Perelman PL, Trifonov VA, Graphodatsky AS. 2012. Chromosomal evolution in rodentia. Heredity (Edinb). 108:4–16.

Rosenberg NA. 2002. The probability of topological concordance of gene trees and species trees. Theor Popul Biol. 61:225–247.

Rowe KC, Achmadi AS, Fabre P-H, Schenk JJ, Steppan SJ, Esselstyn JA. 2019. Oceanic islands of wallacea as a source for dispersal and diversification of murine rodents. Journal of Biogeography. 46:2752–2768.

Roycroft E, Achmadi A, Callahan CM, Esselstyn JA, Good JM, Moussalli A, Rowe KC. 2021. Molecular evolution of ecological specialisation: Genomic insights from the diversification of murine rodents. Genome Biol Evol. 13.

Roycroft EJ, Moussalli A, Rowe KC. 2020. Phylogenomics uncovers confidence and conflict in the rapid radiation of australo-papuan rodents. Syst Biol. 69:431–444.

Sarver BA, Keeble S, Cosart T, Tucker PK, Dean MD, Good JM. 2017. Phylogenomic insights into mouse evolution using a pseudoreference approach. Genome Biol Evol. 9:726–739.

Schenk JJ, Rowe KC, Steppan SJ. 2013. Ecological opportunity and incumbency in the diversification of repeated continental colonizations by muroid rodents. Syst Biol. 62:837–864.

Schliep KP. 2011. Phangorn: Phylogenetic analysis in r. Bioinformatics. 27:592–593.

Serizawa K, Suzuki H, Tsuchiya K. 2000. A phylogenetic view on species radiation in apodemus inferred from variation of nuclear and mitochondrial genes. Biochem Genet. 38:27–40.

Shifman S, Bell JT, Copley RR, Taylor MS, Williams RW, Mott R, Flint J. 2006. A high-resolution single nucleotide polymorphism genetic map of the mouse genome. PLoS Biol. 4:e395.

Smagulova F, Gregoretti IV, Brick K, Khil P, Camerini-Otero RD, Petukhova GV. 2011. Genome-wide analysis reveals novel molecular features of mouse recombination hotspots. Nature. 472:375–378. Repeatmasker open-4.0 [Internet]. 2013-2015. Available from http://www.repeatmasker.org.

Smith JM, Haigh J. 1974. The hitch-hiking effect of a favourable gene. Genet Res. 23:23–35.

Smith MD, Wertheim JO, Weaver S, Murrell B, Scheffler K, Kosakovsky Pond SL. 2015a. Less is more: An adaptive branch-site random effects model for efficient detection of episodic diversifying selection. Mol Biol Evol. 32:1342–1353.

Smith SA, Brown JW, Walker JF. 2018. So many genes, so little time: A practical approach to divergence-time estimation in the genomic era. PLoS One. 13:e0197433.

Smith SA, Moore MJ, Brown JW, Yang Y. 2015b. Analysis of phylogenomic datasets reveals conflict, concordance, and gene duplications with examples from animals and plants. BMC Evol Biol. 15:150.

Stanyon R, Yang F, Cavagna P, O’Brien P, Bagga M, Ferguson-Smith M, Wienberg J. 1999. Animal cytogenetics and comparative mapping-reciprocal chromosome painting shows that genomic rearrangement between rat and mouse proceeds ten times faster than between humans and cats. Cytogenetics and Cell Genetics. 84:150–155.

Steppan SJ, Adkins RM, Spinks PQ, Hale C. 2005. Multigene phylogeny of the old world mice, murinae, reveals distinct geographic lineages and the declining utility of mitochondrial genes compared to nuclear genes. Mol Phylogenet Evol. 37:370–388.

Steppan SJ, Schenk JJ. 2017. Muroid rodent phylogenetics: 900-species tree reveals increasing diversification rates. PLoS One. 12:e0183070.

Sun C, Huang J, Wang Y, Zhao X, Su L, Thomas GWC, Zhao M, Zhang X, Jungreis I, Kellis M, et al. 2021. Genus-wide characterization of bumblebee genomes provides insights into their evolution and variation in ecological and behavioral traits. Mol Biol Evol. 38:486–501.

Suzuki H, Shimada T, Terashima M, Tsuchiya K, Aplin K. 2004. Temporal, spatial, and ecological modes of evolution of eurasian mus based on mitochondrial and nuclear gene sequences. Mol Phylogenet Evol. 33:626–646.

Tange O. 2018. Gnu parallel.

To TH, Jung M, Lycett S, Gascuel O. 2016. Fast dating using least-squares criteria and algorithms. Syst Biol. 65:82–97.

Treaster S, Deelen J, Daane JM, Murabito J, Karasik D, Harris MP. 2023. Convergent genomics of longevity in rockfishes highlights the genetics of human life span variation. Sci Adv. 9:eadd2743.

van der Valk T, Pecnerova P, Diez-Del-Molino D, Bergstrom A, Oppenheimer J, Hartmann S, Xenikoudakis G, Thomas JA, Dehasque M, Saglican E, et al. 2021. Million-year-old DNA sheds light on the genomic history of mammoths. Nature. 591:265–269.

Vanderpool D, Minh BQ, Lanfear R, Hughes D, Murali S, Harris RA, Raveendran M, Muzny DM, Hibbins MS, Williamson RJ, et al. 2020. Primate phylogenomics uncovers multiple rapid radiations and ancient interspecific introgression. PLoS Biol. 18:e3000954.

Wang LG, Lam TT, Xu S, Dai Z, Zhou L, Feng T, Guo P, Dunn CW, Jones BR, Bradley T, et al. 2020. Treeio: An r package for phylogenetic tree input and output with richly annotated and associated data. Mol Biol Evol. 37:599–603.

White MA, Ane C, Dewey CN, Larget BR, Payseur BA. 2009. Fine-scale phylogenetic discordance across the house mouse genome. PLoS Genet. 5:e1000729.

Yalcin B, Wong K, Agam A, Goodson M, Keane TM, Gan X, Nellaker C, Goodstadt L, Nicod J, Bhomra A, et al. 2011. Sequence-based characterization of structural variation in the mouse genome. Nature. 477:326–329.

Yang Z. 2007. Paml 4: Phylogenetic analysis by maximum likelihood. Mol Biol Evol. 24:1586–1591.

Yekutieli D, Benjamini Y. 1999. Resampling-based false discovery rate controlling multiple test procedures for correlated test statistics. Journal of Statistical Planning and Inference. 82:171–196.

Yu G. 2020. Using ggtree to visualize data on tree-like structures. Curr Protoc Bioinformatics. 69:e96.

Yu G, Smith DK, Zhu H, Guan Y, Lam TT-Y. 2017. Ggtree: An r package for visualization and annotation of phylogenetic trees with their covariates and other associated data. Methods in Ecology and Evolution. 8:28–36.

Zhang C, Rabiee M, Sayyari E, Mirarab S. 2018. Astral-iii: Polynomial time species tree reconstruction from partially resolved gene trees. BMC Bioinformatics. 19:153.

