## Supplemental material for "The genomic landscape, causes, and consequences of extensive phylogenomic discordance in Old World mice and rats"

*On UCEs as loci for phylogenetic inference*

Ultra-conserved elements (UCEs) have proven increasingly popular and useful as loci for phylogenomic analyses across a range of taxonomic levels and time scales (Crawford et al. 2012; McCormack et al. 2012; Faircloth et al. 2013; McCormack et al. 2013; Jarvis et al. 2014; Smith et al. 2014; Blaimer et al. 2015; Meiklejohn et al. 2016; Alexander et al. 2017; Burress et al. 2018; Bossert et al. 2019; Oliveros et al. 2019; Quattrini et al. 2020; Alda et al. 2021; McLean et al. 2022). UCEs are typically collected using bait capture and targeted enrichment approaches, harvested from whole genomes as we have done, or a combination of the two. This is driven by both their ready application to non-model organisms and museum specimens (Bi et al. 2013; Faircloth 2017; Zhang et al. 2019), their possession of desirable characters for use in phylogenetic reconstruction (Faircloth et al. 2012; McCormack et al. 2012), and their subsequent ability to provide resolutions to previously intractable nodes (Faircloth et al. 2013; Blaimer et al. 2015; Alda et al. 2021). However, both the function of UCEs and the influence of selection upon them remain poorly understood (Bejerano et al. 2004; Katzman et al. 2007). Rates of molecular evolution may also vary within UCE loci (Tagliacollo and Lanfear 2018) and phylogenetic reconstructions using UCEs (and other phylogenomic markers) can be confounded by gene tree discordance driven by processes such as incomplete lineage sorting and introgression (Jeffroy et al. 2006; Degnan and Rosenberg 2009; Meiklejohn et al. 2016; Chan et al. 2020; Alda et al. 2021).

While UCEs are generally treated as non-coding and independent, they may not be evenly distributed throughout the genome (McCole et al. 2018) and multiple UCEs may overlap the same gene (Van Dam et al. 2021). Merging UCEs that overlap genes can increase topological support but has limited impact on the topology recovered (Van Dam et al. 2021). When available, filtering by phylogenetic signal and noise (Gilbert et al. 2018) can also lead to better supported estimates of tree topology, as can allele phasing of SNPs derived from UCE loci (Andermann et al. 2019; McLean et al. 2022). However, in assessing that support, we notice in our reconstructions and as has been observed elsewhere with multiple phylogenomic markers (Reddy et al. 2017; Chan et al. 2020; Minh et al. 2020; Vanderpool et al. 2020), bootstraps are a poor indicator of support for the correct topology when using phylogenomic data.

UCEs have recently seen increased use in dating analyses and best practices continue to be developed (Blaimer et al. 2015; Branstetter et al. 2017; Bossert et al. 2019; Oliveros et al. 2019; Quattrini et al. 2020). Critically, the scale of UCE datasets can make popular methods of divergence time estimation computationally intractable, while gene tree discordance and substitution rate heterogeneity complicate the selection of appropriate models (Van Dam et al. 2017; Tagliacollo and Lanfear 2018). One strategy to overcome these issues is the selection of a set of loci with model-appropriate properties including clock-like behavior and low gene tree discordance, as described by the SortaDate software package (Smith et al. 2018). Such gene filtering approaches have been suggested as best practice (Walker et al. 2019) and SortaDate specifically has been used in a range of phylogenomic analyses across multiple taxonomic groups (Lind et al. 2019; Del Cortona et al. 2020; Quattrini et al. 2020; Shee et al. 2020; Koenen et al. 2021), though some researchers have found gene filtering has a limited impact on estimated divergence times but increases the associated variance (Oliveros et al. 2019; McGowen et al. 2020).

For dating strategies, least-squares purports vastly reduced computation time with similar accuracy (To et al. 2016) to comparable dating methods implemented in software such as BEAST (Drummond et al. 2006) and r8s (Sanderson 2003). It appears to be robust to topological error (To et al. 2016) and substitution rate heterozygosity between lineages (To et al. 2016; Tong et al. 2018). While developed for dating of rapidly evolving viruses (To et al. 2016), it has seen use in eukaryotes (Yue et al. 2017; Anijalg et al. 2018; Brüniche–Olsen et al. 2018; Tong et al. 2018; Thomas et al. 2019). Given the known rapid evolution of substitution rate in rodents (Douzery et al. 2003), our concerns with tree topology, and its tractability, it is likely appropriate for our dataset of filtered UCE loci.

*Divergence time discussion*

When estimating divergence times on our inferred species trees, the Eumuroidea root (A) is placed at 22.62 Ma, which is concordant with the reconstruction of Schenk et al. (2013). The range between minimum and maximum reconstructed age is wide, overlapping both the maximum first appearance and estimated divergence time for the clade as described by Steppan and Schenk (2017). The estimated date for Muridae (B) of 21.30 Ma likewise has a wide range, and appears to be approximately 0.5 (Schenk et al. 2013), 5 (Chevret and Dobigny 2005), or 8 (Steppan and Schenk 2017) million years older than other estimates. It aligns well with the estimated age of the clade recovered by Aghová et al. (2018) and, with some variance for the dating method used, the supermatrix derived mammalian tree of Meredith et al. (2011) but, as with other nodes, is much younger than was estimated by Hedges et al. (2015). The time of separation of Otomyini and Arvicanthi (Fig. 1, node N) is in general agreement with Lecompte et al. (2008), Schenk et al. (2013), and Aghová et al. (2018) at 8.20 Ma. As with Steppan and Schenk (2017), we reconstruct the origins of Arvicanthini (Fig. 1, node O, 6,56 Ma), Praomyini (node I, 4.82 Ma), and Murini (node K, 6.24 Ma) as approximately 0.5-1 Ma younger than previously determined (Lecompte et al. 2008; Schenk et al. 2013; Aghova et al. 2018), though these older estimates overlap our confidence intervals and Nicolas et al. (2021) have recently estimated an even earlier diversification for Praomyini (7.1 Ma). Despite using a comparatively young calibration range for the *Rattus* group (node Q) our results suggest the clade is possibly even younger given it is reconstructed at the minimum end of the assigned calibration (2.40 Ma), though this may better reflect an internal divergence within *Rattus* sensu lato than the origin of the group. Within the Murini (node K), our divergence of 3.35 Ma for *M. caroli* (node L) followed by the separation of *M. musculus* and *M. spretus* (node O, 1.278 Ma) is very similar to estimates by Suzuki et al. (2004).

*Species tree summary*

Phylogenomic datasets represent a wealth of opportunity to better understand taxonomic relationships, but their size and complexity can make them challenging to use and prone to introducing error (Young & Gillung, 2020; Zhang et al, 2019). As our analyses are based on the available whole genomes of murid rodents, we are inevitably limited in our taxon sampling. Coalescent methods such as ASTRAL-III (Zhang et al., 2018) appear to be resilient to analyses on a small number of taxa (Song et al., 2012; Xi et al., 2015), though adding taxa to a dataset is well understood to improve phylogenetic resolution in general (see Bravo et al., 2019 and citations therein for a discussion). In contrast, adding more genomic data rather than taxa can, counterintuitively, lead to increased support for erroneous topologies (Kumar et al., 2012; Roycroft et al., 2019). It remains to be seen therefore the degree to which the topologies we recover are the result of taxon sampling versus underlying properties of the data, although the extent of gene tree discordance we observe remains striking. Ultimately, choice of data and the models used may matter more than taxon sampling (Reddy et al., 2017), though knowing which data to choose a priori remains an unsolved problem.

Here, we have leveraged the resources of the model organisms the house mouse (*Mus musculus*) and brown rat (*Rattus norvegicus*) along with new genomes from 8 closely related species and eight previously sequenced rodent genomes to understand the systematics of murine rodents and causes and consequences of phylogenetic discordance along the murine genome. Our genomes begin to fill the gap in sampling of murine rodents which, despite their outstanding species diversity, have relatively few whole genomes sequenced and help to place these important model systems in an evolutionary context and provide us with the resources to study the landscape of phylogenetic discordance along the chromosome.

#### Supplemental Figures

#### Figure S1

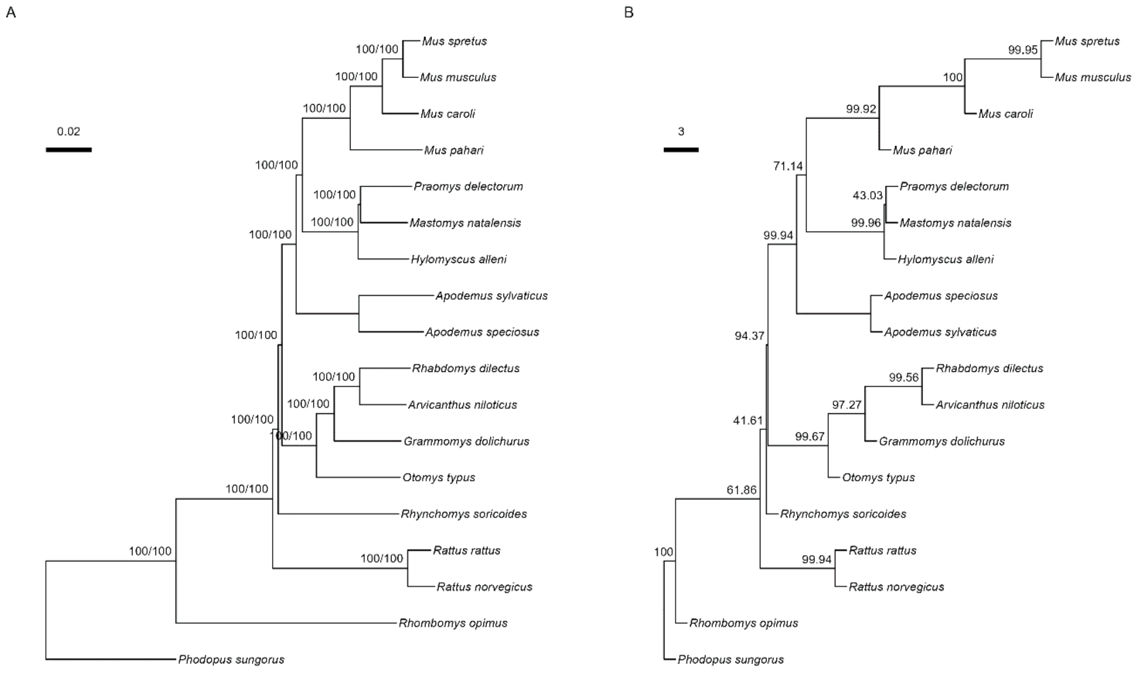


**Figure S1**: Species trees inferred from 2,632 UCE loci. A: Species tree as estimated from concatenation of all loci by IQ-TREE2, with branches showing SH-aLRT/UFBootstrap supports. B: ASTRAL-III species tree based on individual gene trees with internal branch lengths in coalescent units such that shorter branches indicate greater discordance. Branch labels indicate quadripartition support. Final normalized quartet score is 0.81, *i.e.*, 81% of quartet trees induced by the gene trees are represented in the species tree.

**Figure S2**

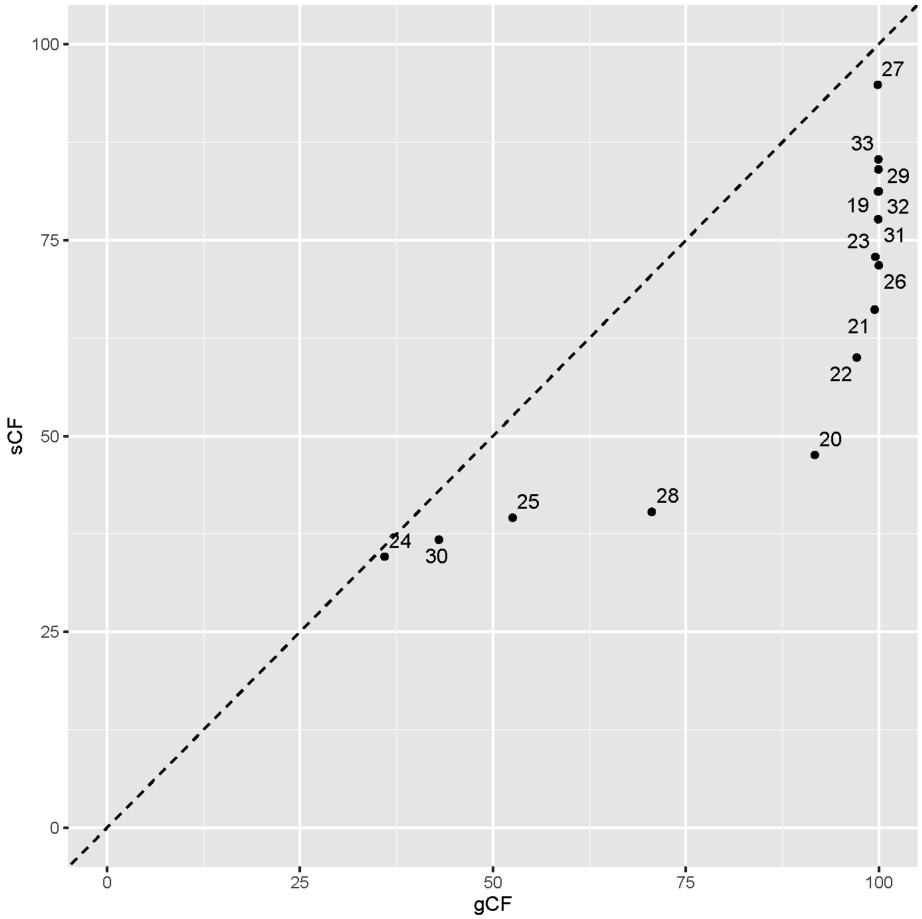


**Figure S2**: Gene-concordance factors (gCF) and site-concordance factors (sCF) for each branch in the concatenated species tree. Note that the lowest possible value for sCF is approximately 30, while gCF can be as low as 0.

**Figure S3**

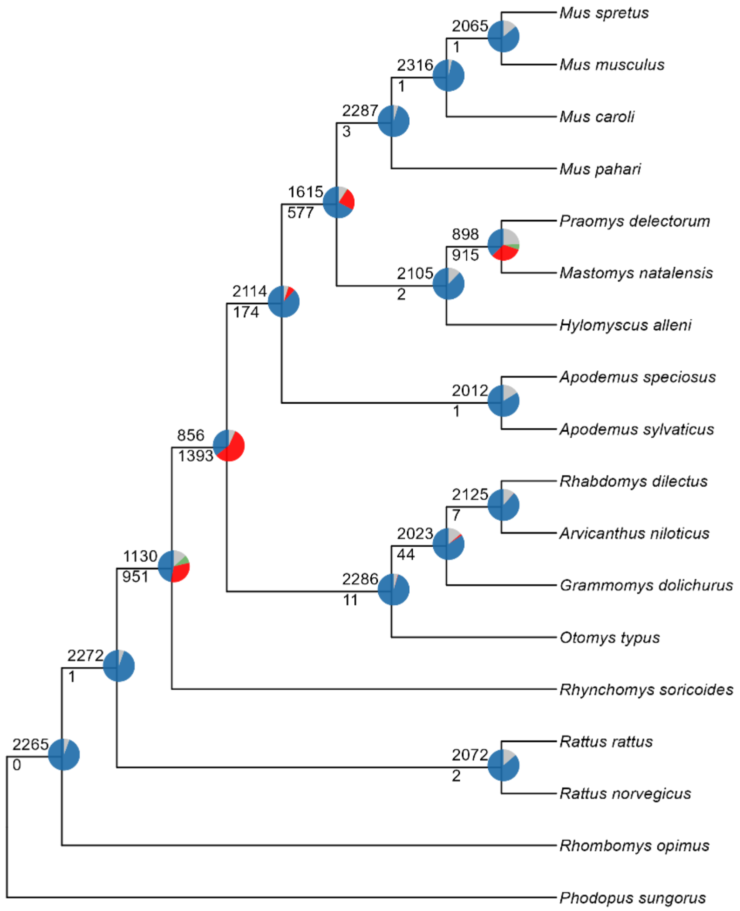


**Figure S3**: Discordance at nodes on the ASTRAL species tree established using the PhyParts package (Smith et al., 2015). The number of gene trees that support the depicted species tree at each node is given above each branch and is represented by the blue portion of the pie chart. The number of gene trees that show a supported conflict with the gene tree is below the branch and is represented in pie charts by green (the most common conflicting partition) and red (all other supported conflicts). The grey section of the pie charts show conflicting gene trees with no support.

**Figure S4**

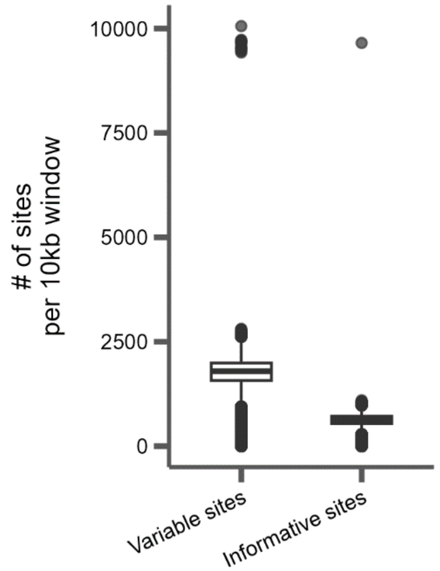


**Figure S4**: The number of variable and informative sites from 165,409 10kb windows from alignments of seven taxa to the mouse reference (mm10) coordinate system. A variable site is defined as any site with more than one allele. An informative site is defined as any site with at least two alleles present in at least two taxa. Average number of variable sites: 1187.8. Average number of informative sites: 401.0.

**Figure S5**

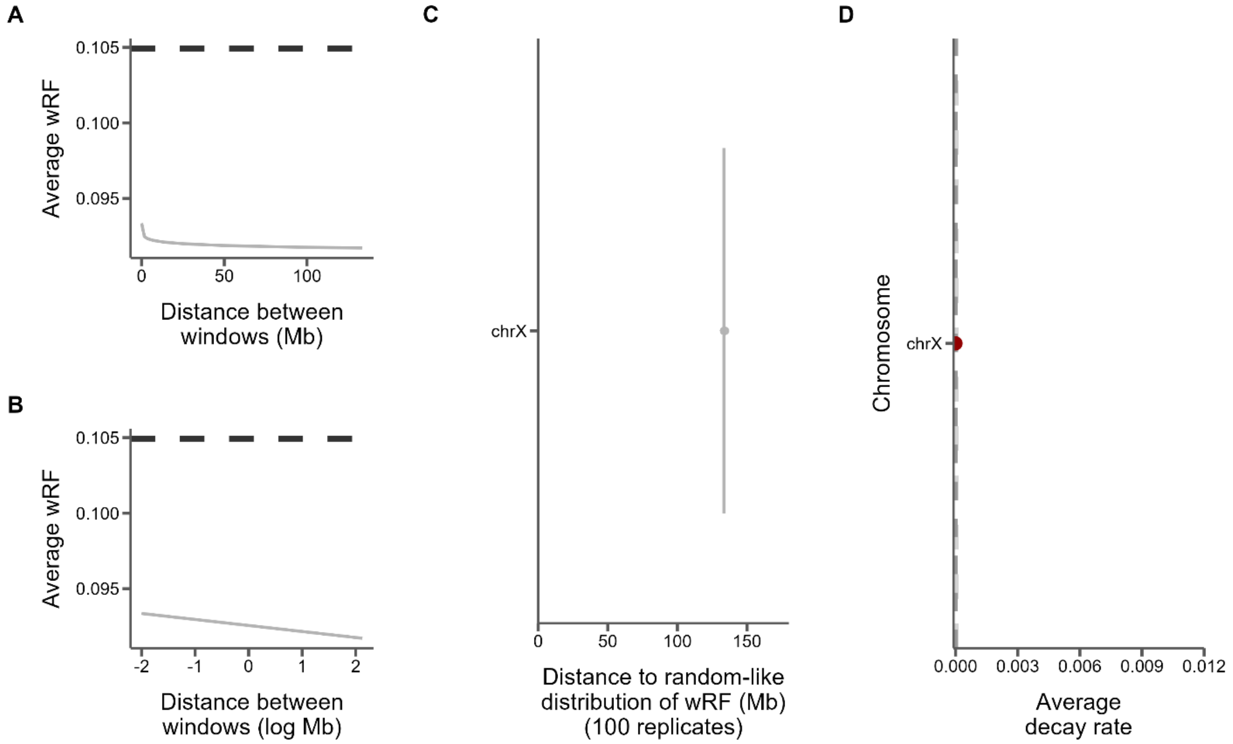


**Figure S5**: Measures of phylogenetic similarity and decay on the X chromosome. A) The log fit to the mean of distributions of tree distances between windows at increasing genomic distance (10kb steps). B) The same, but on a log scale with a linear fit. C) The genomic distance between windows at which tree distance becomes random for 100 replicates of random window selection. D) The slopes of the correlation between genomic distance and tree distance from panel B represent the rate at which tree similarity decays across the chromosome.

**Figure S6**

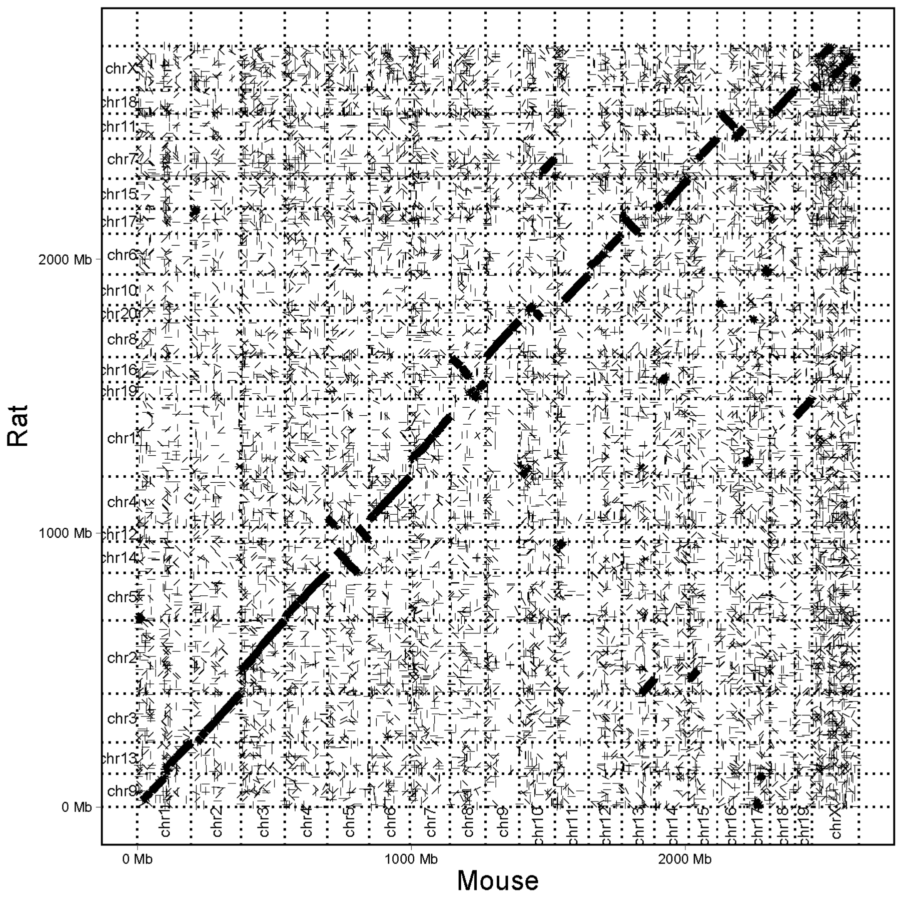


**Figure S6**: Dotplot of whole genome alignment between mouse (mm10) and rat (rnor6) genomes.

**Figure S7**

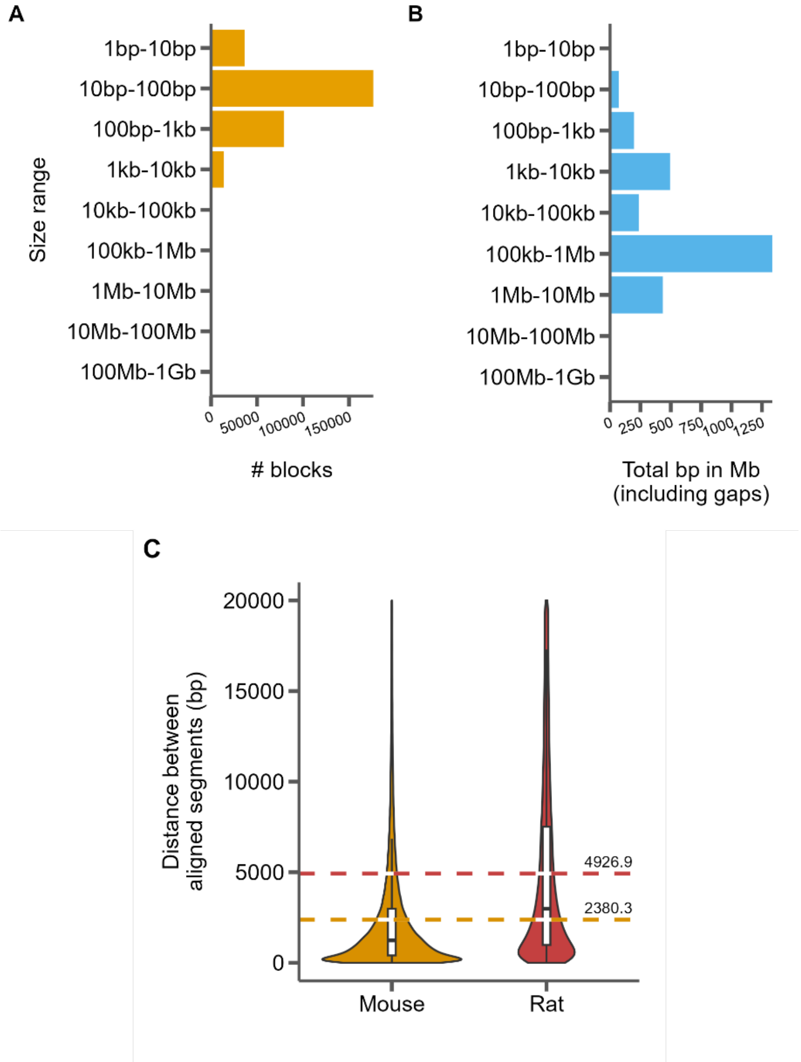


**Figure S7**: Summary of the whole genome alignment between mouse and rat. A) The distribution of aligned block sizes. B) The distribution of the total number of aligned bases for each range of block sizes. C) The distribution of inter-block distances relative to mouse (left) and rat (coordinates) for all blocks shorter than 20 kb. Dashed lines represent average distance between two alignment blocks and are labeled with that average.

**Figure S8**

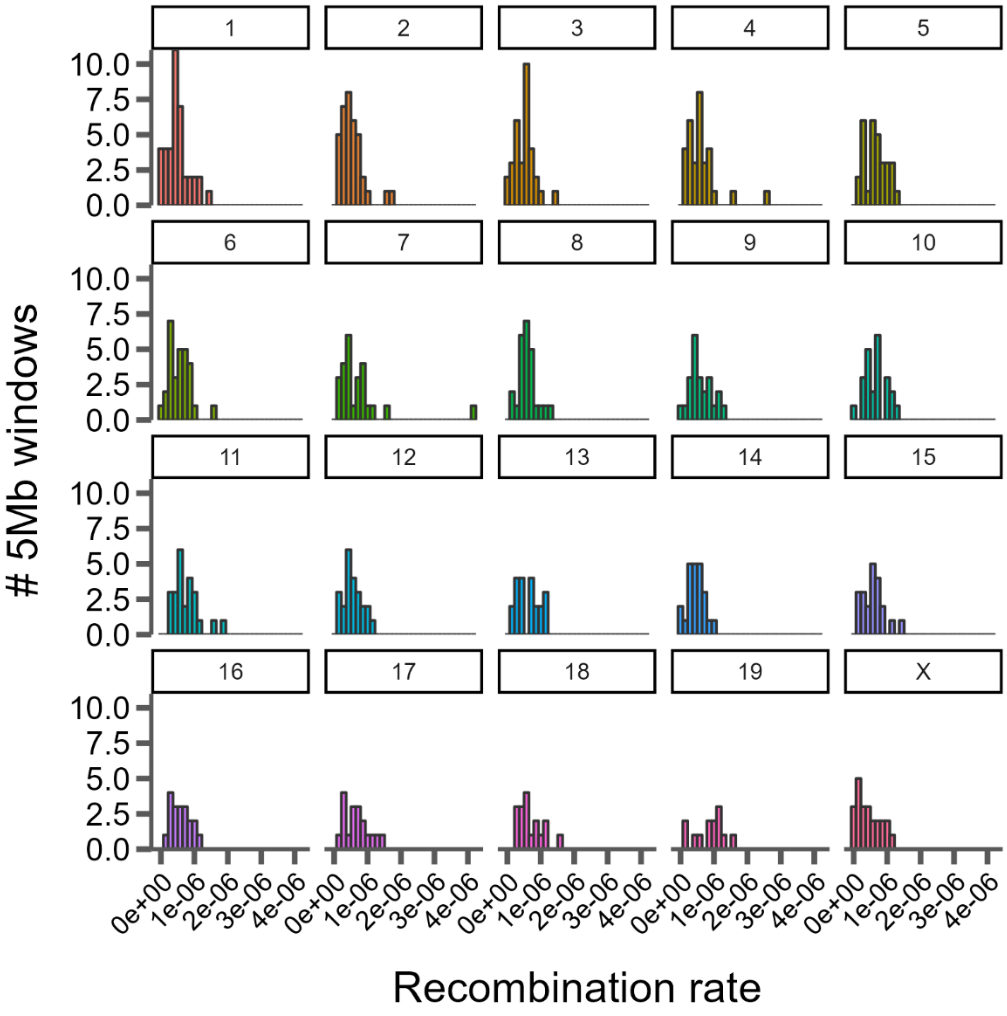


**Figure S8**: Distributions of recombination rate across 19 mouse autosomes and the X chromosome estimated in 5Mb windows. Each line segment represents a single aligned block between the two genomes.

### Figure S9

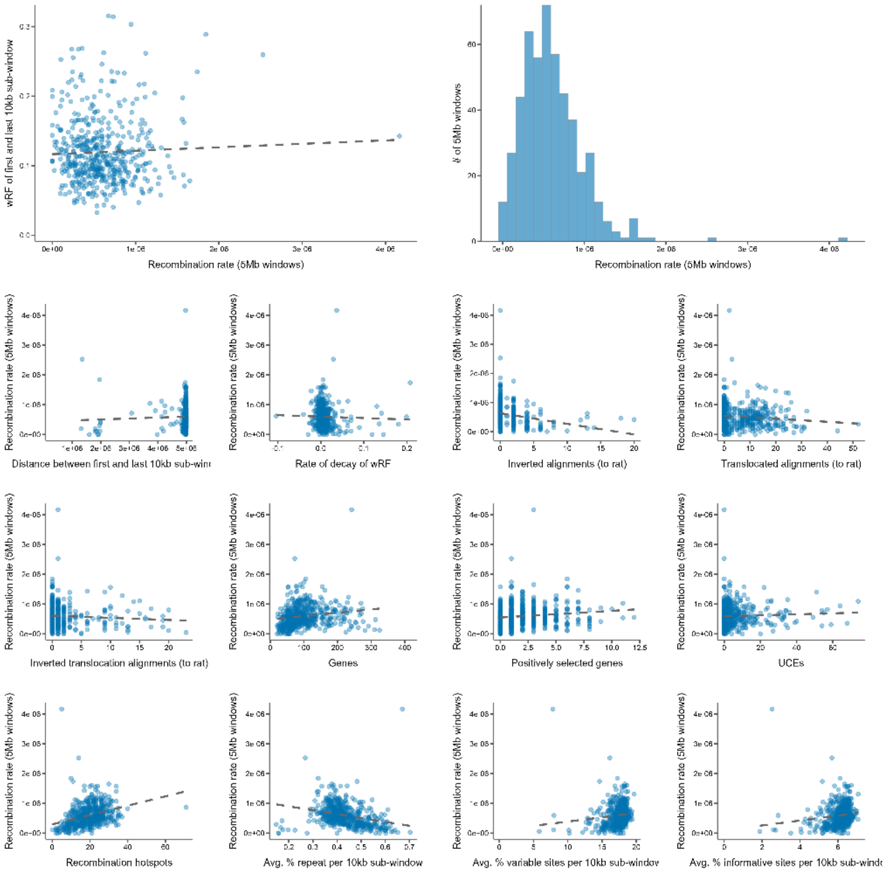


**Figure S9**: Various correlations between recombination rate, phylogenetic similarity, genomic features, and structural variation (from mouse to rat).

**Supplemental Tables**

**Table S1: The estimated age of each node in Figure 1. Nodes with an asterisk (*) indicate those used in calibration, as in Table 2. Node C was used as the fixed calibration (see Methods: Divergence time estimation). Minimum and maximum ages are the lowest and highest values respectively at each node across the three estimated time trees.**

| Node | Estimated Age (mya) | Min/Max Age (mya) |
|---|---|---|
| A | 22.62 | 19.39/25.93 |
| B | 21.30 | 18.43/24.89 |
| C* | 13.08 | 12.10/14.05 |
| D | 12.12 | 11.24 /12.94 |
| E | 11.67 | 10.51/12.83 |
| F | 10.82 | 9.57/12.15 |
| G* | 5.85 | 5.30/7.20 |
| H | 10.18 | 8.71/11.63 |
| I | 4.82 | 3.57/6.00 |
| J | 4.45 | 3.20/5.63 |
| K* | 6.24 | 5.30/7.20 |
| L | 3.35 | 2.62/4.21 |
| M | 1.37 | 0.95/1.87 |
| N | 8.20 | 6.58/10.72 |
| O | 6.56 | 5.05/8.65 |
| P | 4.33 | 3.19/5.60 |
| Q* | 2.40 | 2.40/3.51 |

#### References to the Supplementary Materials


Aghova T, Kimura Y, Bryja J, Dobigny G, Granjon L, Kergoat GJ. 2018. Fossils know it best: Using a new set of fossil calibrations to improve the temporal phylogenetic framework of murid rodents (rodentia: Muridae). *Mol Phylogenet Evol*. 128:98-111.

Alda F, Ludt WB, Elias DJ, McMahan CD, Chakrabarty P. 2021. Comparing ultraconserved elements and exons for phylogenomic analyses of middle american cichlids: When data agree to disagree. *Genome Biol Evol*. 13.

Alexander AM, Su YC, Oliveros CH, Olson KV, Travers SL, Brown RM. 2017. Genomic data reveals potential for hybridization, introgression, and incomplete lineage sorting to confound phylogenetic relationships in an adaptive radiation of narrow-mouth frogs. *Evolution*. 71:475-488.

Andermann T, Fernandes AM, Olsson U, Topel M, Pfeil B, Oxelman B, Aleixo A, Faircloth BC, Antonelli A. 2019. Allele phasing greatly improves the phylogenetic utility of ultraconserved elements. *Syst Biol*. 68:32-46.

Anijalg P, Ho SYW, Davison J, Keis M, Tammeleht E, Bobowik K, Tumanov IL, Saveljev AP, Lyapunova EA, Vorobiev AA, et al. 2018. Large-scale migrations of brown bears in eurasia and to north america during the late pleistocene. *Journal of Biogeography*. 45:394-405.

Bejerano G, Pheasant M, Makunin I, Stephen S, Kent WJ, Mattick JS, Haussler D. 2004. Ultraconserved elements in the human genome. *Science*. 304:1321-1325.

Bi K, Linderoth T, Vanderpool D, Good JM, Nielsen R, Moritz C. 2013. Unlocking the vault: Next-generation museum population genomics. *Mol Ecol*. 22:6018-6032.

Blaimer BB, Brady SG, Schultz TR, Lloyd MW, Fisher BL, Ward PS. 2015. Phylogenomic methods outperform traditional multi-locus approaches in resolving deep evolutionary history: A case study of formicine ants. *BMC Evol Biol*. 15:271.

Bossert S, Murray EA, Almeida EAB, Brady SG, Blaimer BB, Danforth BN. 2019. Combining transcriptomes and ultraconserved elements to illuminate the phylogeny of apidae. *Mol Phylogenet Evol*. 130:121-131.

Branstetter MG, Danforth BN, Pitts JP, Faircloth BC, Ward PS, Buffington ML, Gates MW, Kula RR, Brady SG. 2017. Phylogenomic insights into the evolution of stinging wasps and the origins of ants and bees. *Current Biology*. 27:1019-1025.

Brüniche–Olsen A, Jones ME, Burridge CP, Murchison EP, Holland BR, Austin JJ. 2018. Ancient DNA tracks the mainland extinction and island survival of the tasmanian devil. *Journal of Biogeography*. 45:963-976.

Burress ED, Alda F, Duarte A, Loureiro M, Armbruster JW, Chakrabarty P. 2018. Phylogenomics of pike cichlids (cichlidae: Crenicichla): The rapid ecological speciation of an incipient species flock. *J Evol Biol*. 31:14-30.

Chan KO, Hutter CR, Wood PL, Jr., Grismer LL, Brown RM. 2020. Target-capture phylogenomics provide insights on gene and species tree discordances in old world treefrogs (anura: Rhacophoridae). *Proc Biol Sci*. 287:20202102.

Chevret P, Dobigny G. 2005. Systematics and evolution of the subfamily gerbillinae (mammalia, rodentia, muridae). *Mol Phylogenet Evol*. 35:674-688.

Crawford NG, Faircloth BC, McCormack JE, Brumfield RT, Winker K, Glenn TC. 2012. More than 1000 ultraconserved elements provide evidence that turtles are the sister group of archosaurs. *Biol Lett*. 8:783-786.

Degnan JH, Rosenberg NA. 2009. Gene tree discordance, phylogenetic inference and the multispecies coalescent. *Trends Ecol Evol*. 24:332-340.

Del Cortona A, Jackson CJ, Bucchini F, Van Bel M, D'Hondt S, Skaloud P, Delwiche CF, Knoll AH, Raven JA, Verbruggen H, et al. 2020. Neoproterozoic origin and multiple transitions to macroscopic growth in green seaweeds. *Proc Natl Acad Sci U S A*. 117:2551-2559.

Douzery EJ, Delsuc F, Stanhope MJ, Huchon D. 2003. Local molecular clocks in three nuclear genes: Divergence times for rodents and other mammals and incompatibility among fossil calibrations. *J Mol Evol*. 57 Suppl 1:S201-213.

Drummond AJ, Ho SY, Phillips MJ, Rambaut A. 2006. Relaxed phylogenetics and dating with confidence. *PLoS Biol*. 4:e88.

Faircloth BC. 2017. Identifying conserved genomic elements and designing universal bait sets to enrich them. *Methods in Ecology and Evolution*. 8:1103-1112.

Faircloth BC, McCormack JE, Crawford NG, Harvey MG, Brumfield RT, Glenn TC. 2012. Ultraconserved elements anchor thousands of genetic markers spanning multiple evolutionary timescales. *Syst Biol*. 61:717-726.

Faircloth BC, Sorenson L, Santini F, Alfaro ME. 2013. A phylogenomic perspective on the radiation of ray-finned fishes based upon targeted sequencing of ultraconserved elements (uces). *PLoS One*. 8:e65923.

Gilbert PS, Wu J, Simon MW, Sinsheimer JS, Alfaro ME. 2018. Filtering nucleotide sites by phylogenetic signal to noise ratio increases confidence in the neoaves phylogeny generated from ultraconserved elements. *Mol Phylogenet Evol*. 126:116-128.

Hedges SB, Marin J, Suleski M, Paymer M, Kumar S. 2015. Tree of life reveals clock-like speciation and diversification. *Mol Biol Evol*. 32:835-845.

Jarvis ED, Mirarab S, Aberer AJ, Li B, Houde P, Li C, Ho SY, Faircloth BC, Nabholz B, Howard JT, et al. 2014. Whole-genome analyses resolve early branches in the tree of life of modern birds. *Science*. 346:1320-1331.

Jeffroy O, Brinkmann H, Delsuc F, Philippe H. 2006. Phylogenomics: The beginning of incongruence? *Trends Genet*. 22:225-231.

Katzman S, Kern AD, Bejerano G, Fewell G, Fulton L, Wilson RK, Salama SR, Haussler D. 2007. Human genome ultraconserved elements are ultraselected. *Science*. 317:915.

Koenen EJM, Ojeda DI, Bakker FT, Wieringa JJ, Kidner C, Hardy OJ, Pennington RT, Herendeen PS, Bruneau A, Hughes CE. 2021. The origin of the legumes is a complex paleopolyploid phylogenomic tangle closely associated with the cretaceous-paleogene (k-pg) mass extinction event. *Syst Biol*. 70:508-526.

Lecompte E, Aplin K, Denys C, Catzeflis F, Chades M, Chevret P. 2008. Phylogeny and biogeography of african murinae based on mitochondrial and nuclear gene sequences, with a new tribal classification of the subfamily. *BMC Evol Biol*. 8:199.

Lind AL, Lai YYY, Mostovoy Y, Holloway AK, Iannucci A, Mak ACY, Fondi M, Orlandini V, Eckalbar WL, Milan M, et al. 2019. Genome of the komodo dragon reveals adaptations in the cardiovascular and chemosensory systems of monitor lizards. *Nat Ecol Evol*. 3:1241-1252.

McCole RB, Erceg J, Saylor W, Wu CT. 2018. Ultraconserved elements occupy specific arenas of three-dimensional mammalian genome organization. *Cell Rep*. 24:479-488.

McCormack JE, Faircloth BC, Crawford NG, Gowaty PA, Brumfield RT, Glenn TC. 2012. Ultraconserved elements are novel phylogenomic markers that resolve placental mammal phylogeny when combined with species-tree analysis. *Genome Res*. 22:746-754.

McCormack JE, Harvey MG, Faircloth BC, Crawford NG, Glenn TC, Brumfield RT. 2013. A phylogeny of birds based on over 1,500 loci collected by target enrichment and high-throughput sequencing. *PLoS One*. 8:e54848.

McGowen MR, Tsagkogeorga G, Alvarez-Carretero S, Dos Reis M, Struebig M, Deaville R, Jepson PD, Jarman S, Polanowski A, Morin PA, et al. 2020. Phylogenomic resolution of the cetacean tree of life using target sequence capture. *Syst Biol*. 69:479-501.
McLean BS, Bell KC, Cook JA. 2022. Snp-based phylogenomic inference in holarctic ground squirrels (urocitellus). *Mol Phylogenet Evol*. 169:107396.
Meiklejohn KA, Faircloth BC, Glenn TC, Kimball RT, Braun EL. 2016. Analysis of a rapid evolutionary radiation using ultraconserved elements: Evidence for a bias in some multispecies coalescent methods. *Syst Biol*. 65:612-627.
Meredith RW, Janecka JE, Gatesy J, Ryder OA, Fisher CA, Teeling EC, Goodbla A, Eizirik E, Simao TL, Stadler T, et al. 2011. Impacts of the cretaceous terrestrial revolution and kpg extinction on mammal diversification. *Science*. 334:521-524.
Minh BQ, Hahn MW, Lanfear R. 2020. New methods to calculate concordance factors for phylogenomic datasets. *Mol Biol Evol*. 37:2727-2733.
Nicolas V, Mikula O, Lavrenchenko LA, Sumbera R, Bartakova V, Bryjova A, Meheretu Y, Verheyen E, Missoup AD, Lemmon AR, et al. 2021. Phylogenomics of african radiation of praomyini (muridae: Murinae) rodents: First fully resolved phylogeny, evolutionary history and delimitation of extant genera. *Mol Phylogenet Evol*. 163:107263.
Oliveros CH, Field DJ, Ksepka DT, Barker FK, Aleixo A, Andersen MJ, Alstrom P, Benz BW, Braun EL, Braun MJ, et al. 2019. Earth history and the passerine superradiation. *Proc Natl Acad Sci U S A*. 116:7916-7925.
Quattrini AM, Rodriguez E, Faircloth BC, Cowman PF, Brugler MR, Farfan GA, Hellberg ME, Kitahara MV, Morrison CL, Paz-Garcia DA, et al. 2020. Palaeoclimate ocean conditions shaped the evolution of corals and their skeletons through deep time. *Nat Ecol Evol*. 4:1531-1538.
Reddy S, Kimball RT, Pandey A, Hosner PA, Braun MJ, Hackett SJ, Han KL, Harshman J, Huddleston CJ, Kingston S, et al. 2017. Why do phylogenomic data sets yield conflicting trees? Data type influences the avian tree of life more than taxon sampling. *Syst Biol*. 66:857-879.
Sanderson MJ. 2003. R8s: Inferring absolute rates of molecular evolution and divergence times in the absence of a molecular clock. *Bioinformatics*. 19:301-302.
Schenk JJ, Rowe KC, Steppan SJ. 2013. Ecological opportunity and incumbency in the diversification of repeated continental colonizations by muroid rodents. *Syst Biol*. 62:837-864.
Shee ZQ, Frodin DG, Camara-Leret R, Pokorny L. 2020. Reconstructing the complex evolutionary history of the papuasian schefflera radiation through herbariomics. *Front Plant Sci*. 11:258.
Smith BT, Harvey MG, Faircloth BC, Glenn TC, Brumfield RT. 2014. Target capture and massively parallel sequencing of ultraconserved elements for comparative studies at shallow evolutionary time scales. *Syst Biol*. 63:83-95.
Smith SA, Brown JW, Walker JF. 2018. So many genes, so little time: A practical approach to divergence-time estimation in the genomic era. *PLoS One*. 13:e0197433.
Steppan SJ, Schenk JJ. 2017. Muroid rodent phylogenetics: 900-species tree reveals increasing diversification rates. *PLoS One*. 12:e0183070.
Suzuki H, Shimada T, Terashima M, Tsuchiya K, Aplin K. 2004. Temporal, spatial, and ecological modes of evolution of eurasian mus based on mitochondrial and nuclear gene sequences. *Mol Phylogenet Evol*. 33:626-646.
Tagliacollo VA, Lanfear R. 2018. Estimating improved partitioning schemes for ultraconserved elements. *Mol Biol Evol*. 35:1798-1811.
Thomas JE, Carvalho GR, Haile J, Rawlence NJ, Martin MD, Ho SY, Sigfusson A, Josefsson VA, Frederiksen M, Linnebjerg JF, et al. 2019. Demographic reconstruction from ancient DNA supports rapid extinction of the great auk. *Elife*. 8.

To TH, Jung M, Lycett S, Gascuel O. 2016. Fast dating using least-squares criteria and algorithms. *Syst Biol*. 65:82-97.

Tong KJ, Duchene DA, Duchene S, Geoghegan JL, Ho SYW. 2018. A comparison of methods for estimating substitution rates from ancient DNA sequence data. *BMC Evol Biol*. 18:70.

Van Dam MH, Henderson JB, Esposito L, Trautwein M. 2021. Genomic characterization and curation of uces improves species tree reconstruction. *Syst Biol*. 70:307-321.

Van Dam MH, Lam AW, Sagata K, Gewa B, Laufa R, Balke M, Faircloth BC, Riedel A. 2017. Ultraconserved elements (uces) resolve the phylogeny of australasian smurf-weevils. *PLoS One*. 12:e0188044.

Vanderpool D, Minh BQ, Lanfear R, Hughes D, Murali S, Harris RA, Raveendran M, Muzny DM, Hibbins MS, Williamson RJ, et al. 2020. Primate phylogenomics uncovers multiple rapid radiations and ancient interspecific introgression. *PLoS Biol*. 18:e3000954.

Walker JF, Walker-Hale N, Vargas OM, Larson DA, Stull GW. 2019. Characterizing gene tree conflict in plastome-inferred phylogenies. *PeerJ*. 7:e7747.

Yue JX, Li J, Aigrain L, Hallin J, Persson K, Oliver K, Bergstrom A, Coupland P, Warringer J, Lagomarsino MC, et al. 2017. Contrasting evolutionary genome dynamics between domesticated and wild yeasts. *Nat Genet*. 49:913-924.

Zhang YM, Williams JL, Lucky A. 2019. Understanding uces: A comprehensive primer on using ultraconserved elements for arthropod phylogenomics. *Insect Systematics and Diversity*. 3.
